# Tracing urban diet shifts into guppy stoichiometry in Trinidadian streams

**DOI:** 10.64898/2026.11.15.766775

**Authors:** Jeferson R. Amaral, Amina Mohamed, Peter C. Searle, Stephanie M. Tran, Jillon Lewis, Amy Deacon, Swanne Gordon, Andrés López-Sepulcre

## Abstract

1. Urbanization can alter dietary stoichiometry by changing the elemental composition of food resources or the mixture of resources consumers assimilate. However, the relative contributions of resource elemental quality and assimilated diet composition to dietary stoichiometry, and whether the resulting dietary differences are reflected in consumer body stoichiometry, remain poorly understood, particularly in omnivores.
2. We sampled adult female Trinidadian guppies (*Poecilia reticulata*) and their food resources from paired high-urbanization (HU) and low-urbanization (LU) sites in three rivers of Trinidad’s Northern Range. Combining hierarchical Bayesian stable-isotope mixing models with resource %C, %N, and %P, we reconstructed dietary stoichiometry, partitioned differences between urbanization levels into contributions from diet composition and resource elemental quality, and compared these with somatic stoichiometry.
3. Under the primary mixing model, aquatic invertebrates contributed less to estimated HU diets (33.7 percentage points lower; 95% credible interval [5.0, 60.1]), whereas detrital contributions tended to be higher. Resource elemental composition showed no consistent differences between urbanization levels.
4. Under this model, reconstructed dietary N:P ratios were lower in HU guppies (by 59.2% [10.0, 81.8]), while C:N ratios tended to be higher, and C:P ratios tended to be lower. Changes in assimilated source mixtures contributed more strongly than resource elemental quality to C:N and N:P differences. All of these contrasts depended on the estimated trophic discrimination factors, weakening under fixed literature values, with the C:N contrast reversing under two of them and the detrital contrast varying in direction.
5. Guppy somatic tissues partly tracked these dietary differences. HU guppies tended to have lower somatic C:P and N:P ratios, in the same direction as their diets but of smaller magnitude. Somatic C:N showed no clear difference, and the dietary C:N contrast was itself uncertain.
6. Our findings suggest that urbanization reshapes guppy dietary stoichiometry mainly through shifts in which resources they assimilate, and that guppies only partially pass these shifts into their bodies. Understanding the stoichiometric consequences of environmental change therefore requires considering not only what resources contain, but also which resources consumers assimilate.

## INTRODUCTION

Food webs link consumer feeding to the movement of carbon (C), nitrogen (N), and phosphorus (P) through ecosystems. The elemental composition, or stoichiometry, of consumer diets shapes how these elements are acquired, retained, and recycled (Sterner & Elser, 2002). Environmental change can alter dietary stoichiometry through two pathways: changes in the stoichiometry of available resources and changes in the relative contributions of those resources to the diet (Mehler et al., 2013; Cross et al., 2003; Zhang et al., 2018). This distinction is especially relevant for omnivores, which integrate multiple trophic pathways and can adjust resource use as environmental conditions change (Remonti et al., 2016; Wootton, 2017). Yet, how changes in resource stoichiometry and dietary composition jointly shape the elemental balance of vertebrate consumers under environmental change remains poorly understood.

Urbanization provides a useful context for examining how changes in resource use and resource stoichiometry may jointly influence consumer dietary stoichiometry. In streams, urban land use can modify riparian canopy cover, nutrient and contaminant inputs, organic matter sources, and prey communities (Carroll & Jackson, 2008; Paul & Meyer, 2001; Roy et al., 2003; Walsh et al., 2005). These environmental changes can alter the relative availability and consumption of different food resources, including algae, detritus, and animal prey (Eitzmann & Paukert, 2010; O’Brien & Wehr, 2010; Peressin et al., 2018; Singer & Battin, 2007). For example, the omnivorous goby *Awaous stamineus* increased its reliance on animal prey in urban Hawaiian streams (Lisi et al., 2018), while some fishes in disturbed Brazilian streams increased their consumption of detritus (Peressin et al., 2018). Urbanization can also affect the elemental composition of food resources, with studies documenting N-enriched leaf litter and lower leaf-litter C:N ratios (Milanovich et al., 2014), as well as increased C, N, and P content in periphyton and changes in fish stoichiometry (Montaña & Schalk, 2018). Thus, urbanization can simultaneously change what consumers eat and the elemental composition of those resources, potentially altering dietary stoichiometry through both pathways.

Previous studies have documented elemental variation across food-web components and used realized diets to estimate consumer-resource elemental imbalances (Lauridsen et al., 2014; Milanovich et al., 2014; Montaña & Schalk, 2018; Singer & Battin, 2007; Tsoi et al., 2011). Work on western mosquitofish (*Gambusia affinis*) has also linked environmentally associated differences in diet composition to consumer body stoichiometry, although without reconstructing the elemental composition of assimilated diets (Moffett et al., 2021). However, the relative contributions of changes in resource stoichiometry and assimilated resource mixtures to dietary stoichiometry remain unclear, as does the extent to which these dietary differences translate into changes in consumer body stoichiometry. Distinguishing these pathways is essential for understanding how environmental change propagates through food webs to influence consumer elemental composition.

The Trinidadian guppy (*Poecilia reticulata*) provides an opportunity to investigate these relationships in an omnivorous consumer that feeds on algae, detritus, and aquatic invertebrates (Zandonà et al., 2011, 2015, 2017). A previous study has shown that body stoichiometry differs substantially among Trinidadian streams (El-Sabaawi et al., 2012). Despite considerable variation in reconstructed dietary C:P ratios among streams, that study found no significant relationship between dietary and body C:P ratios. In contrast, experimental manipulation of dietary stoichiometry demonstrated that P-enriched diets increase whole-body phosphorus and reduce C:P and N:P ratios, without affecting C (Dalton et al., 2017). More recently, Zandonà et al. (2024) found that tissue N increased with the proportion of invertebrates consumed across natural populations, whereas somatic phosphorus was not associated with diet quality. Together, these findings suggest that dietary stoichiometry can influence guppy stoichiometry, but that its effects under natural conditions remain difficult to predict. Urbanization offers an opportunity to investigate this relationship because urban guppy populations exhibit differences in resource use, trophic interactions, and resource-acquisition traits (Amaral et al., 2021; de Carvalho et al., 2019; Marques et al., 2021; Santana Marques et al., 2020), including evolved differences in gut length (Rigdon et al., 2026). Whether urbanization-associated changes in resource composition and feeding combine to alter dietary stoichiometry, and whether these changes are reflected in guppy body stoichiometry, remains unresolved.

Using paired high-urbanization (HU) and low-urbanization (LU) sites in three rivers of Trinidad’s Northern Range, we tested two non-exclusive hypotheses for how urbanization alters the stoichiometry of guppy diets (Figure 1). Under the Resource Quality Hypothesis, urbanization changes the elemental composition of individual food resources, altering their nutritional quality and consequently dietary stoichiometry, independently of changes in the relative contribution of different resources. For example, nutrient enrichment may increase resource N or P content relative to C, reducing dietary C:N and C:P ratios (Granitto et al., 2022; Kohler et al., 2012; Stelzer & Lamberti, 2001). However, responses may differ among resources (Bumpers et al., 2023; Singer & Battin, 2007). Under the Diet Composition Hypothesis, urbanization changes the relative contributions of algae, detritus, and animal prey to guppy diets, altering dietary stoichiometry through shifts in resource consumption, independently of changes in the elemental composition of individual resources. For example, greater reliance on relatively N-poor basal resources rather than N-rich animal prey could increase dietary C:N ratios, with effects on C:P and N:P depending on the elemental composition of the resources being substituted.

**Figure 1.**
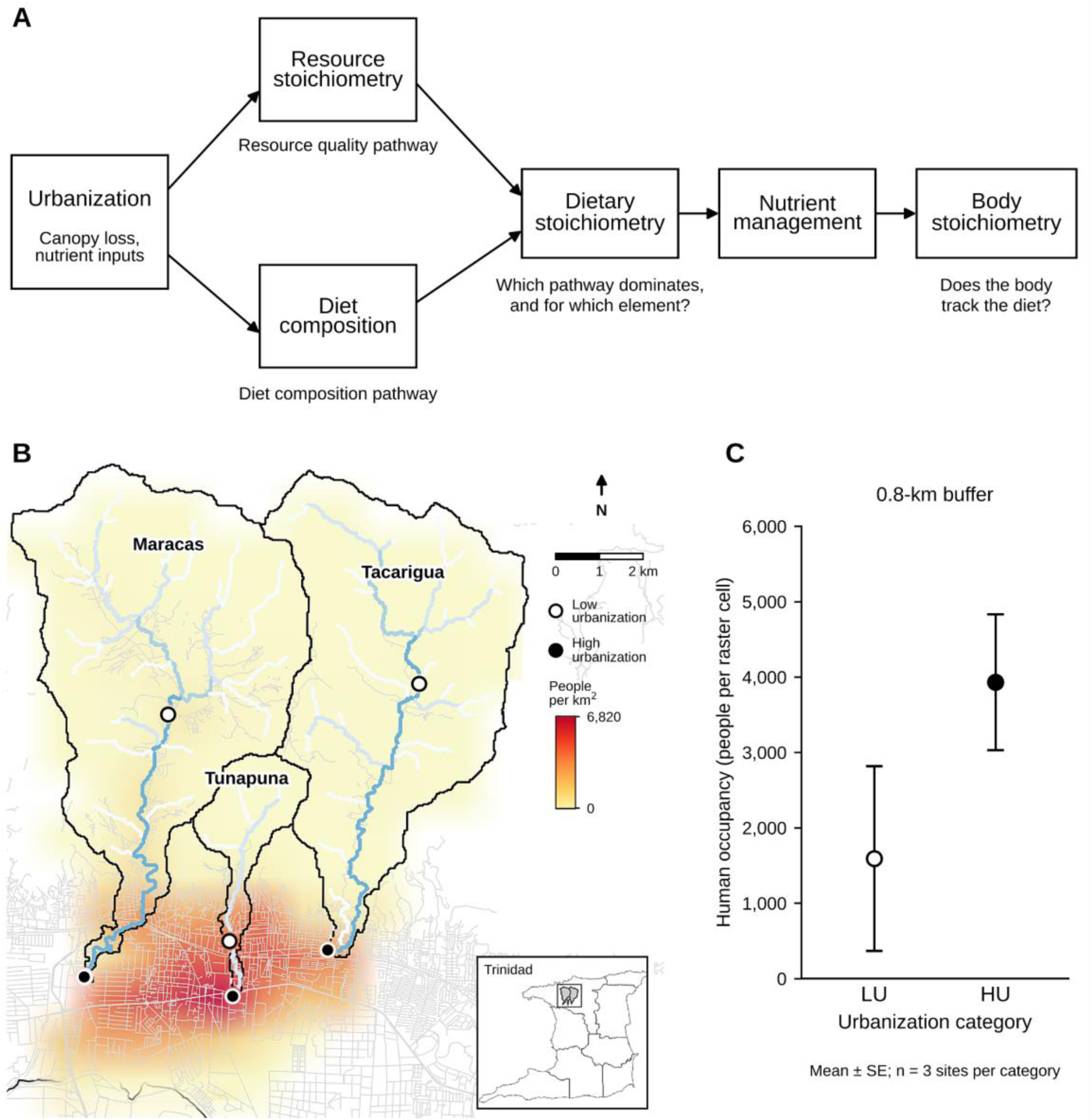
Conceptual framework, study sites, and urbanization context in Trinidad. (A) Two pathways through which urbanization may influence dietary stoichiometry: changes in the elemental composition of available resources (resource quality pathway) and changes in assimilated diet composition (diet composition pathway). Dietary stoichiometry may subsequently influence body stoichiometry through consumer nutrient management. (B) Location of paired high-urbanization (HU; filled circles) and low-urbanization (LU; open circles) sampling sites in the Maracas, Tacarigua, and Tunapuna rivers, Trinidad. Grayscale shading represents estimated human population density (people km⁻^2^), and dark gray lines indicate the main river channels. The inset shows the location of the study area within Trinidad. (C) Raster-derived human occupancy, expressed as mean people per raster cell within a 0.8-km buffer around each site. Points represent model-estimated means for each urbanization category from a mixed model with River, Site, and year random intercepts, and error bars indicating ±1 SE. Open and filled circles denote LU and HU sites, respectively.

To evaluate these hypotheses, we combined stable-isotope estimates of assimilated resource mixtures with measurements of resource elemental composition and partitioned HU–LU differences in reconstructed dietary C:N, C:P, and N:P. We then examined whether differences in reconstructed dietary stoichiometry were reflected in guppy somatic stoichiometry. This allowed us to assess whether changes in dietary elemental composition translate into changes in consumer body composition, or whether physiological regulation buffers these effects.

## MATERIALS AND METHODS

### Study Sites

Before field sampling, we identified candidate sites along three rivers in Trinidad’s Northern Range (Maracas/St. Joseph, Tunapuna, and Caura/Tacarigua) using human population density estimates from the WorldPop raster (1 km resolution) (WorldPop, 2020) as an initial proxy for urbanization. Sites were provisionally classified as high-urbanization (HU; > 1,000 ind. km^-2^) and low-urbanization (LU; < 500 ind. km^-2^).

In March of 2023, we surveyed candidate sites to assess accessibility, evidence of human disturbance (i.e., discharge pipes, nearby housing, and channelization), and guppy presence. We then selected one HU and one LU site within each river (Figure 1). To minimize confounding effects of variation in predation pressure, we avoided upstream reaches where predation pressure naturally declines (Deacon et al., 2018) and selected sites with comparable predator communities across urbanization categories (Supplementary Material, *Fish Community Analysis*).

We characterized environmental conditions using watershed-scale land cover, impervious surface cover, and road density, together with local measurements of canopy cover, channel structure, and water quality. We subsequently validated our HU–LU classification using human occupancy estimates from the LandScan Mosaic raster (Adams et al., 2025) across multiple spatial buffers (Supplementary Material, “*Environmental characterization, urbanization assessment, and principal component analysis*”).

### Resource and Fish Sampling

To characterize trophic pathways across HU and LU sites, we sampled epilithon, filamentous algae, aquatic macroinvertebrates, and detrital resources, including coarse particulate organic matter (CPOM), fine particulate organic matter (FPOM), and sewage inputs from pipelines when present. We also collected snails at each site to establish site-specific δ^15^N baselines, and riparian grasses as a terrestrial isotopic reference. All samples were analyzed for %C, %N, %P, δ^13^C, and δ^15^N (Supplementary Material, “*Resource collection and processing*”).

At each site, we collected 30 adult female guppies using butterfly nets for elemental and stable-isotope analyses. Fish were euthanized using MS-222, frozen, and later dissected to remove the gastrointestinal tract and gonads. We examined gut contents from 38 remaining fish across five sites (no gut-content data were available for Tunapuna LU because individuals were prioritized for stable-isotope and elemental analyses) under a dissecting microscope and recorded the presence of each food item (Table S19). Food-item occurrence frequencies were used to inform the prior distribution of dietary source contributions in the isotope mixing model (see below). The remaining somatic tissue was dried at 50 °C to a constant mass and homogenized with a mortar and pestle.

### Elemental and Isotope Analyses

To quantify elemental carbon and nitrogen concentrations and stable isotope signatures in resources and fish tissues, we measured %C and %N and δ^13^C and δ^15^N at the Cornell Isotope Laboratory using elemental analysis coupled to isotope-ratio mass spectrometry. We determined phosphorus concentrations in approximately 2-mg aliquots of dried resource and fish tissue. Samples were combusted at 550 °C for 1 h, and the resulting ash was digested in 10 mL of ultrapure water and 2 mL of 2 N HCl at 105 °C for 2 h. Phosphorus concentrations were then determined colorimetrically using the ascorbic acid method (Murphy & Riley, 1962).

### Scales of Inference

The scales of inference and replication for all analyses are summarized in Table 1. Urbanization was classified at the site level, with one high-urbanization (HU) and one low-urbanization (LU) site paired within each of three rivers. Thus, the three river pairs provided the primary spatial replication for comparisons between urbanization categories, whereas individual fish and resource observations represented subsamples within sites. To account for this sampling structure, models testing urbanization category (Origin) or site-level environmental predictors included random intercepts for River and Site (defined by River × Origin), with additional random effects where appropriate. Frequentist analyses were used to characterize environmental and community patterns, including associations between environmental gradients and resource stoichiometry. Bayesian models were used to estimate assimilated dietary composition, reconstruct dietary stoichiometry, and evaluate somatic elemental composition and stoichiometry.

**Table 1.** Scales of inference and replication. All analyses compare 3 HU and 3 LU sites paired within 3 rivers, with river and site as random intercepts.

| Analysis | Scale at which urbanization is applied | Subsamples within sites |
|---|---|---|
| Site characterization (land cover, water quality, occupancy, PCA) | Site-level classification; occupancy also has year as a random effect | 4 site-year occupancy records per site (24 total) |
| Resource elemental composition and stoichiometry | Site-level fixed effect, fitted per resource type | 292 resource records |
| Assimilated diet and dietary stoichiometry | Site-level fixed effect in the isotope mixing model, with dry mass as a covariate; decomposition from paired posterior draws | 30 fish per site (180 total) |
| Somatic dry mass | Site-level fixed effect, with dry mass as a covariate for stoichiometry | 30 fish per site (180 total) |

### Diet Composition

We estimated the assimilated dietary composition of guppies using hierarchical Bayesian stable isotope mixing models implemented in JAGS (Plummer, 2003; Semmens et al., 2009; Ward et al., 2010). The analysis included 180 guppies and 292 dietary resource observations from the six sites. We considered four resource categories: detritus, epilithon, filamentous algae, and aquatic invertebrates. Source isotope signatures were estimated hierarchically, accounting for variation among rivers, sites, and resource subcategories. We modeled dietary proportions using an isometric log-ratio transformation, with urbanization category (Origin) as a fixed effect, river and site as random effects, and standardized log-transformed somatic dry mass as a covariate (Supplementary Material, “*Hierarchical Bayesian stable isotope mixing models*”).

We lipid-normalized fish δ^13^C values (Post et al., 2007) and centered consumer and resource δ^15^N values relative to site-specific snail baselines (Post, 2002). The mixing model accounted for source %C and %N and estimated carbon and nitrogen trophic discrimination factors (TDFs) jointly with resource isotope signatures and dietary proportions. TDFs were assigned literature-informed normal priors (carbon: 0.40 ± 1.30‰; nitrogen: 2.54 ± 1.27‰, mean ± SD; Post, 2002; Vanderklift & Ponsard, 2003). Resource and consumer isotope observations were modeled jointly, propagating uncertainty in source isotope signatures into dietary estimates. A single gut-content-informed Dirichlet prior, balanced across urbanization categories, informed baseline dietary composition (Supplementary Material, “*Modeling consumer dietary composition*”; Table S2).

We evaluated sensitivity to TDF assumptions using three additional models with fixed literature-derived discrimination factors. We also evaluated sensitivity to alternative dietary priors (uniform Dirichlet), urbanization coding (symmetric rather than reference-level), and assumptions about site-level dietary heterogeneity (wider prior on the site-effect standard deviation; Supplementary Material, “*Trophic discrimination estimation and sensitivity analyses*”, “*Sensitivity to urbanization-prior parameterization*”, and “*Sensitivity to dietary prior specifications*”). Complete model specifications, priors, diagnostics, and sensitivity analyses are provided in the Supplementary Material. Model convergence and adequacy were evaluated using rank-normalized *R^*, effective sample sizes, and posterior predictive checks. We also evaluated the influence of consumer isotope observations on estimated resource signatures using a source-only hierarchical model (Supplementary Material, “*Source-only diagnostic and model evaluation*” and “*Trophic discrimination estimates and source-model diagnostics*”). HU−LU dietary contrasts were estimated within each river at the reference somatic mass (29.68 mg, the pooled geometric mean) and averaged equally across the three river pairs. We ran four chains of 1,000,000 iterations, discarding the first 500,000 as burn-in and thinning by 100, which retained 20,000 posterior draws. We report posterior medians, 95% credible intervals (CrI), and directional posterior probabilities.

### Estimating and Decomposing Dietary Stoichiometry

To reconstruct the elemental composition of assimilated diets, we combined posterior estimates of dietary source proportions from the hierarchical isotope mixing model with estimates of resource %C, %N, and %P. We fitted 12 separate Bayesian models, one for each resource-element combination, including urbanization category (Origin) as a fixed effect and river and site as random intercepts. Detritus models additionally included organic matter subtypes (CPOM, FPOM, and sewage-derived material) as a fixed effect, whereas invertebrate models included taxon as a random intercept. Elemental concentrations were modeled using a lognormal distribution, except detrital %C, which was modeled using beta regression to constrain predictions to the physically possible range. We used empirically calibrated, weakly informative priors, with a zero-centered prior for the urbanization effect (Supplementary Material, “*Prior specification and sensitivity of resource elemental composition models*”).

For each resource category, we estimated elemental concentrations separately for HU and LU sites from the fitted Bayesian models, setting river and site random-effect deviations to zero to obtain urbanization-category estimates. For aquatic invertebrates, predictions retained taxon-specific random effects and were averaged equally across the 26 modeled taxa. This standardized resource composition across taxa without assuming that sampling frequencies reflected prey availability. Detrital predictions retained the fixed effects of organic matter subtype, with CPOM, FPOM, and sewage-derived material weighted equally.

We summarized dietary source contributions using all 20,000 retained posterior draws from the hierarchical mixing model. For dietary stoichiometry reconstruction, we randomly selected 10,000 joint dietary posterior draws and paired them with 10,000 draws from the independently fitted resource elemental models, using conditional arithmetic mean elemental concentrations. For each draw, dietary elemental concentrations were calculated as the sum of source proportions multiplied by their corresponding elemental concentrations. We then calculated molar C:N, C:P, and N:P ratios using atomic masses. The reconstruction assumes proportional assimilation of elements across resources and does not account for correlations among resource elemental concentrations. Within each draw, we also expressed the contribution of each source to dietary C, N, and P, to identify the pathways underlying urbanization-associated differences in elemental supply.

To partition HU–LU differences in dietary stoichiometry, we reconstructed four combinations of dietary source proportions and resource elemental composition within each posterior draw: HU proportions with HU resources, HU proportions with LU resources, LU proportions with HU resources, LU proportions with LU resources. The diet-composition component represents the average effect of changing dietary source proportions while holding resource elemental composition constant, whereas the resource-quality component represents the average effect of changing resource elemental composition while holding dietary proportions constant. Each component was averaged across the two possible reference conditions. Contrasts were calculated on the log-ratio scale, allowing the two components to sum exactly to the total HU–LU difference. Contrasts were back-transformed into relative percentage differences and summarized using posterior medians, 95% credible intervals, and directional posterior probabilities. We also calculated the posterior probability that the absolute diet-composition contribution exceeded that of resource quality.

Finally, we evaluated the sensitivity of dietary reconstruction to alternative resource elemental assumptions by using conditional medians instead of arithmetic means, site-balanced occurrence weights for macroinvertebrate taxa and detrital subtypes, and excluding sewage-derived material from detrital elemental predictions. We also examined hypothetical scenarios with greater sewage contributions to detritus in HU than LU populations. To evaluate whether dietary stoichiometry depended on discrimination assumptions, we repeated the reconstruction using dietary posteriors from the three fixed-TDF models. These analyses tested the robustness of dietary estimates to assumptions about resource composition and isotope discrimination (Supplementary Material, “*Sensitivity of dietary reconstruction to resource elemental prediction scale*”, “*Sensitivity to detrital subtype weighting*”, “*Sensitivity to hypothetical sewage enrichment*”, and “*Sensitivity to macroinvertebrate taxon weighting*”).

### Somatic Stoichiometry

Because body size may differ between urbanization categories, we first compared log somatic dry mass between origins using a linear mixed model with River and Site random intercepts. We then evaluated whether HU–LU differences in reconstructed dietary stoichiometry were accompanied by differences in somatic stoichiometry. For that, we fitted separate Bayesian Gaussian mixed-effects models for somatic C:N, C:P, and N:P molar ratios, as well as %C, %N, and %P. Somatic responses were log-transformed, and models were fitted in brms with weakly informative priors. Models included Origin and standardized log somatic dry mass as fixed effects, with River and Site as random intercepts. We estimated HU–LU contrasts at a common somatic dry mass and summarized them using posterior medians, 95% credible intervals, and directional posterior probabilities. Contrasts were expressed as relative percentage differences, with additional percentage-point differences calculated for elemental percentages. Because dietary and somatic models were fitted separately, we paired draws from their marginal posterior distributions to compare dietary and somatic responses descriptively. Using the same paired draws, we calculated a diet-to-body index for each molar ratio as the reconstructed dietary ratio divided by the corresponding mass-adjusted somatic ratio. Values above 1 indicate a diet richer in the first element than the body. The index describes differences between dietary and somatic elemental ratios, rather than homeostatic regulation, and its HU–LU contrast indicates whether urbanization-associated differences were greater in diet or body stoichiometry.

### Software

All statistical analyses were conducted in R (R Core Team, 2024). We organized and summarized data using the tidyverse suite of packages (Wickham et al., 2019). We implemented the hierarchical Bayesian stable isotope mixing models in JAGS using the rjags interface (Plummer, 2003) and modeled resource elemental composition and somatic stoichiometry using brms, which fits models in Stan (Bürkner, 2017). We fitted linear mixed-effects models using lme4 (Bates et al., 2015) and lmerTest (Kuznetsova et al., 2017), and beta-distributed generalised linear mixed-effects models using glmmTMB (Brooks et al., 2017). We summarized multivariate environmental variation using principal component analysis (PCA) implemented with FactoMineR and factoextra (Lê et al., 2008), and we analyzed community composition with vegan (Oksanen et al., 2007). We produced figures with ggplot2 (Wickham, 2016).

## RESULTS

### Site Characteristics

Raster-derived human occupancy supported our a priori classification of sites into high- and low-urbanization categories (Figure 1). Compared with paired low-urbanization (LU) sites, high-urbanization (HU) sites generally had greater human occupancy, development, and impervious surface cover, but lower tree and canopy cover (Table 2). In contrast, differences in dissolved nutrients and other water-quality variables were less consistent among rivers. Further environmental characterization is provided in the Supplementary Material (*Environmental characterization, urbanization assessment, and principal component analysis*).

**Table 2.** Environmental characteristics of low and high urbanization habitats.

| Scale | Environmental variable | Unit | Low urbanization | High urbanization |
|---|---|---|---|---|
| **Watershed** | Tree cover | % | 95.63 ± 2.86 | 90.26 ± 5.68 |
|  | Impervious surface | % | 2.04 ± 2.69 | 6.89 ± 5.97 |
|  | Built-up cover | % | 2.69 ± 2.82 | 7.30 ± 6.38 |
|  | Grassland cover | % | 1.59 ± 0.89 | 2.33 ± 1.13 |
|  | Road density | km km<sup>-2</sup> | 1.23 ± 0.95 | 2.60 ± 1.64 |
| **Local buffer** | Raster-derived human occupancy | mean people cell<sup>-1</sup> | 1591.417 ± 2483.525 | 3930.771 ± 1784.692 |
| **In-stream** | Canopy cover | % | 65.27 ± 4.29 | 16.39 ± 15.03 |
|  | Benthic area | m² | 179.46 ± 49.76 | 154.79 ± 42.92 |
|  | Temperature | °C | 26.70 ± 0.95 | 29.17 ± 2.01 |
|  | Dissolved oxygen | % | 79.20 ± 24.45 | 90.73 ± 36.26 |
|  | pH | pH units | 7.48 ± 0.10 | 7.69 ± 0.25 |
|  | Conductivity | µS cm<sup>-1</sup> | 447.00 ± 199.23 | 531.00 ± 67.13 |
|  | Dissolved inorganic N | mmol L<sup>-1</sup> | 0.0124 ± 0.0086 | 0.0207 ± 0.0092 |
|  | Dissolved P | mmol L<sup>-1</sup> | 0.0028 ± 0.0008 | 0.0087 ± 0.0095 |
|  | Dissolved N:P | molar ratio | 4.58 ± 2.88 | 3.79 ± 1.94 |
|  | Fecal coliforms | counts 100 mL<sup>-1</sup> | 551.33 ± 908.46 | 563.33 ± 897.78 |

*Environmental variables measured at the watershed, local-buffer, and in-stream scales across three rivers. Values represent means ± standard deviations (SD) across three sites per urbanization category. Raster-derived human occupancy represents the mean estimated ambient population count per raster cell within a 0.8-km buffer around each site. Dissolved inorganic nitrogen was calculated from ammonium, nitrite, and nitrate concentrations. Environmental variables are presented descriptively to characterize differences between urbanization categories.*

### Resource stoichiometry did not differ consistently with urbanization

We found no clear evidence of differences in elemental concentrations or molar ratios between HU and LU sites across the four resource categories (all 24 posterior 95% credible intervals included zero; Figure 2). Associations between resource stoichiometry and environmental gradients are presented in the Supplementary Material (“*Exploratory associations between environmental variables and resource stoichiometry*”; Table S1; Figure S4).

**Figure 2.**
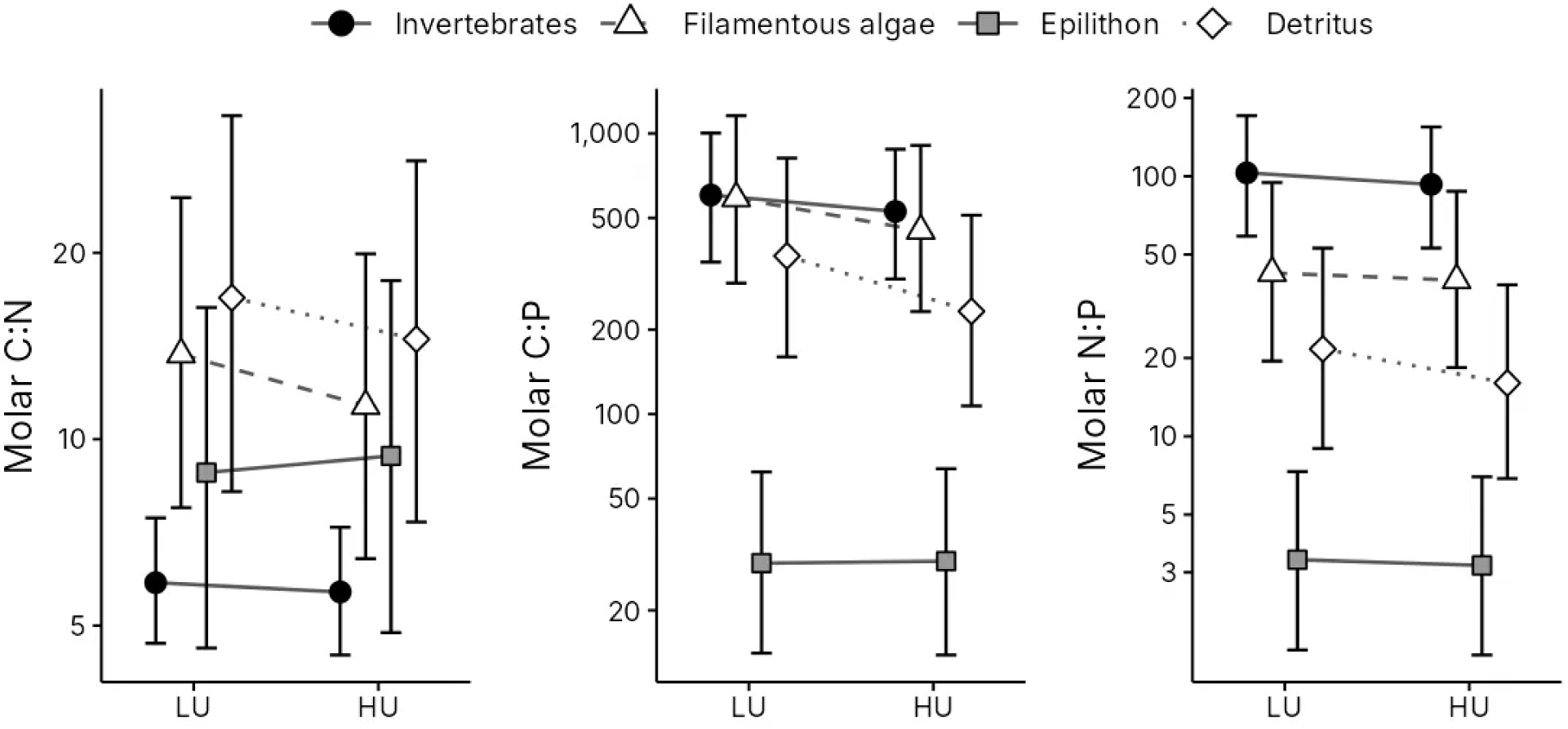
Stoichiometry of dietary resources in low- and high-urbanization streams. Model-predicted molar C:N, C:P and N:P ratios of aquatic invertebrates (filled circles), filamentous algae (open triangles), epilithon (grey squares) and detritus (open diamonds) at low-urbanization (LU) and high-urbanization (HU) sites. Points show origin-level posterior medians, error bars show 95% credible intervals, and lines connect the LU and HU estimates for each resource. Resource ratios did not differ clearly between urbanization origins: all 95% credible intervals for the HU vs LU differences overlapped zero. Predictions exclude river- and site-level random-effect deviations. The y axes are on a logarithmic scale.

### Urban guppies assimilated less invertebrate prey

Under our primary mixing model, aquatic invertebrates contributed substantially less to the assimilated diets of high-urbanization (HU) guppies than to those of low-urbanization (LU) guppies (posterior medians: 5.6% and 41.3%, respectively; Figure 3). The paired HU–LU contrast was −33.7 percentage points (95% CrI [−60.1, −5.0]; *Pr*[HU < LU] = 0.991), with negative median contrasts across all three rivers. Detritus showed the opposite, although less certain pattern, contributing 54.2% of assimilated diet in HU populations compared with 19.3% in LU populations (HU–LU: +33.0 percentage points, 95% CrI [−4.6, +64.2]; *Pr*[HU > LU] = 0.954). Neither epilithon nor filamentous algae showed clear directional differences between urbanization categories.

**Figure 3.**
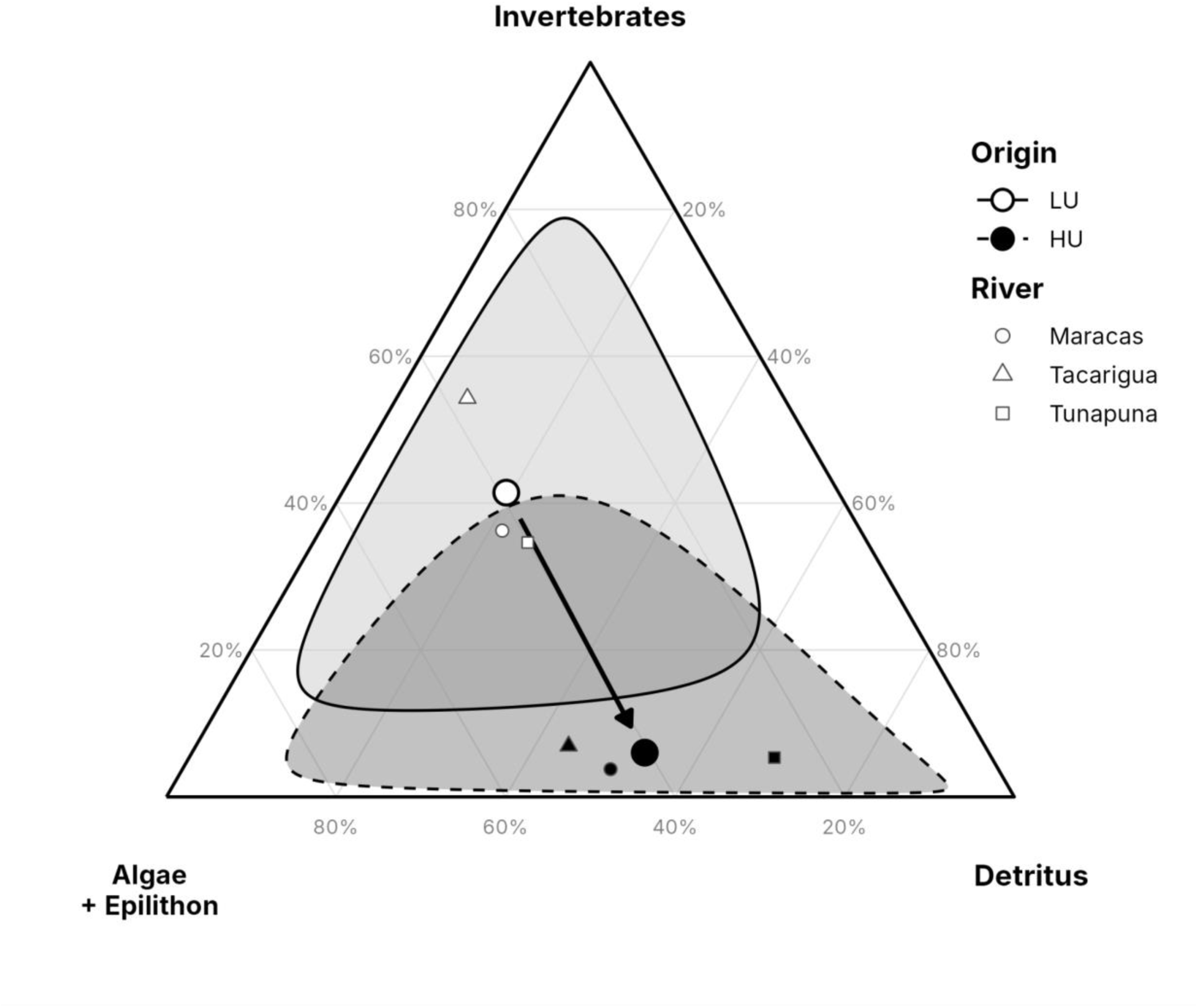
Assimilated diet of guppies from low (LU, white) and high (HU, black) urbanization sites, estimated with a stable isotope mixing model. Large symbols show the average of the three rivers, with shaded 95% credible regions, and the arrow shows the change from LU to HU. Small symbols show individual sites. Position in the triangle gives the proportion of each food source; algae refers to filamentous algae and epilithon, combined for visualization. Estimates are for a fish of reference body mass. Estimated dietary contributions and paired HU–LU contrasts are reported in Supplementary Tables S3 and S4, respectively.

These dietary contrasts were sensitive to assumptions about trophic discrimination factors (TDFs). Because TDFs have not been experimentally calibrated for guppy somatic tissue, our primary mixing model estimated carbon and nitrogen TDFs jointly with diet. The estimated carbon TDF (2.82‰, 95% CrI [2.20, 3.33]) was substantially higher than its literature-informed prior mean (0.40‰), whereas the nitrogen TDF (3.00‰ [2.72, 3.41]) remained close to its prior mean (2.54‰). Under all three alternative models using fixed literature TDFs, the median invertebrate contrast remained negative but was weaker, with 95% credible intervals overlapping zero; detrital contrasts varied in direction (Supplementary Material, “Sensitivity to trophic discrimination factors”; Table S5). Sensitivity analyses using symmetric urbanization coding and a wider prior on dietary site-level heterogeneity produced similar directional contrasts (Supplementary Material, “*Sensitivity to urbanization-prior parameterization*” and “*Sensitivity to dietary prior specifications*”).

We next examined how differences in inferred dietary composition translated into the relative contributions of food sources to assimilated C, N, and P. Under the primary mixing model, aquatic invertebrates contributed less to all three elements in HU than LU guppies, with HU–LU differences of −49.4 percentage points for C (95% CrI [−73.4, −11.4]; *Pr*[HU<LU] = 0.996),−53.5 for N ([−78.9, −14.3]; *Pr* = 0.997), and −41.2 for P ([−68.0, −7.9]; *Pr* = 0.994). Conversely, detritus contributed more to assimilated C (+38.1 percentage points, 95% CrI [+0.8, +67.0]; *Pr*[HU>LU] = 0.978), N (+34.3 [+1.5, +66.2]; *Pr* = 0.982), and P (+40.2 [+0.6, +70.7]; *Pr* = 0.977) in HU populations. Filamentous algae and epilithon showed no clear directional differences in their elemental contributions.

### Diet composition was the main contributor to differences in dietary stoichiometry

Under the primary model, inferred dietary N:P ratios were 59.2% lower in HU than LU guppies (95% CrI [−81.8, −10.0]; *Pr*[HU < LU] = 0.987; Figure 4). Dietary C:P ratios also tended to be lower (−41.3%, 95% CrI [−69.7, +10.5]; *Pr* = 0.954), whereas C:N ratios tended to be higher (+42.7%, 95% CrI [−9.0, +132.6]; *Pr* = 0.940), although the credible intervals for both contrasts included zero. The C:N contrast was particularly sensitive to TDF assumptions, reversing direction under two of the three fixed-TDF specifications (Table S6).

**Figure 4.**
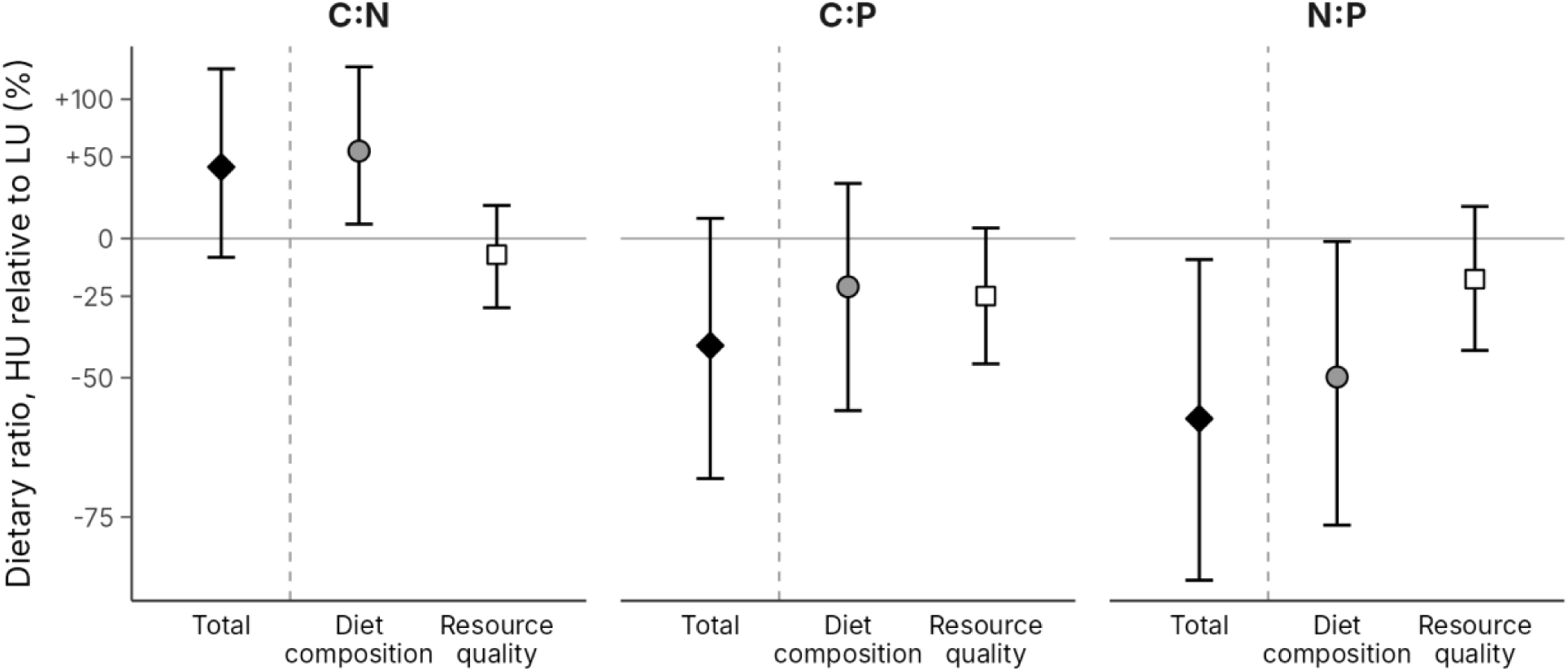
Sources of the difference in dietary stoichiometry between guppies from high urbanization (HU) and low urbanization (LU) sites. For each molar ratio, the total difference in reconstructed dietary stoichiometry (black diamonds, left of the dashed line) is split into two components: the shift in the proportions of resources assimilated (diet composition, grey circles) and differences in the elemental content of the resources themselves (resource quality, open squares). Differences were calculated within each paired river and posterior draw, then averaged across the three rivers. Points are posterior medians and error bars are 95% credible intervals. The y-axis uses a logarithmic transformation with tick labels expressed as HU–LU percentage changes. Diet-composition and resource-quality contributions sum to the total on the log-ratio scale, but not after back-transformation to percentages. The horizontal line indicates no difference. Component estimates are given in Table 3.

**Table 3.** Diet, somatic stoichiometry, and diet-to-body indices of guppies from LU and HU sites.

|  | LU | HU | HU vs LU difference | Pr(direction) |
|---|---|---|---|---|
| **Assimilated diet (%)** |  |  | *percentage points* |  |
| Aquatic invertebrates | 41.3 [12.8, 67.5] | 5.6 [0.8, 26.2] | −33.7 [−60.1, −5.0] | 0.991 |
| Detritus | 19.3 [4.5, 43.4] | 54.2 [11.9, 83.5] | +33.0 [−4.6, +64.2] | 0.954 |
| Filamentous algae | 13.9 [5.1, 37.6] | 18.2 [4.6, 42.0] | +2.9 [−15.4, +25.0] | 0.631 |
| Epilithon | 21.4 [1.4, 52.3] | 15.9 [0.5, 60.4] | −3.0 [−32.3, +33.4] | 0.613 |
| **Dietary stoichiometry (molar)** |  |  | *% change* |  |
| C:N | 7.4 [5.9, 11.1] | 10.8 [7.2, 17.3] | +42.7 [−9.0, +132.6] | 0.940 |
| Diet composition |  |  | +54.5 [+7.4, +134.9] |  |
| Resource quality |  |  | −7.7 [−29.2, +17.8] |  |
| C:P | 445.8 [254.3, 685.2] | 260.2 [127.6, 468.1] | −41.3 [−69.7, +10.5] | 0.954 |
| Diet composition |  |  | −21.4 [−57.6, +31.5] |  |
| Resource quality |  |  | −24.9 [−46.4, +5.3] |  |
| N:P | 59.2 [29.6, 97.4] | 23.7 [10.9, 48.2] | −59.2 [−81.8, −10.0] | 0.987 |
| Diet composition |  |  | −49.8 [−76.0, −1.5] |  |
| Resource quality |  |  | −18.3 [−42.7, +17.3] |  |
| **Somatic stoichiometry (molar)** |  |  | *% change* |  |
| C:N | 4.6 [4.1, 5.2] | 4.4 [3.8, 5.2] | −4.2 [−12.4, +5.8] | 0.898 |
| C:P | 29.9 [24.3, 36.4] | 24.9 [18.6, 33.6] | −16.6 [−32.8, +3.8] | 0.957 |
| N:P | 6.5 [5.2, 8.2] | 5.6 [4.1, 7.9] | −12.8 [−29.8, +10.7] | 0.910 |
| **Diet-to-body index** |  |  | *% change* |  |
| C:N | 1.6 [1.2, 2.4] | 2.4 [1.6, 4.0] | +48.6 [−5.9, +145.1] | 0.957 |
| C:P | 14.9 [8.3, 24.1] | 10.4 [4.8, 20.2] | −29.6 [−64.9, +37.8] | 0.855 |
| N:P | 9.1 [4.4, 15.8] | 4.2 [1.8, 9.1] | −53.4 [−79.8, +6.5] | 0.965 |

*Note. Posterior medians [95% credible intervals] from the primary mixing model. Pr(direction) is the posterior probability of the sign of the median difference. Differences were averaged across the three river pairs; the diet composition and resource quality components sum to the total on the log scale. Pr(\|composition\| > \|quality\|) was 0.935 for C:N, 0.472 for C:P, and 0.908 for N:P. The diet-to-body index is the dietary ratio divided by the somatic ratio. Somatic estimates and indices refer to a common dry mass. Results under fixed discrimination factors are in Tables S5 and S6.*

Decomposing the dietary stoichiometric differences into contributions from diet composition and resource quality revealed stronger effects of diet composition on C:N and N:P (Figure 4). Changes in diet composition favored higher dietary C:N ratios in HU guppies (+54.5%, 95% CrI [+7.4, +134.9]) and lower N:P ratios (−49.8% [−76.0, −1.5]), whereas the corresponding contributions of resource quality were smaller, with credible intervals overlapping zero. The posterior probability that the absolute contribution of diet composition exceeded that of resource quality was 0.935 for C:N and 0.908 for N:P. In contrast, diet composition and resource quality contributed similarly to the lower dietary C:P ratios in HU guppies (diet composition: −21.4%, 95% CrI [−57.6, +31.5]; resource quality: −24.9% [−46.4, +5.3]), but neither component was clearly dominant (*Pr*[|composition| > |quality|] = 0.472).

### Somatic elemental ratios tended to be lower in urban guppies

HU guppies were generally smaller than LU guppies (somatic dry mass: 26.4 ± 17.4 versus 48.0 ± 31.8 mg, mean ± SD). Geometric mean dry mass was lower at the HU site in all three rivers (by 17% in Maracas, 40% in Tacarigua, and 57% in Tunapuna), although the overall contrast was uncertain (−41%, 95% CI [−74, +35]; t_2_ = −2.71, p = 0.11; Supplementary Material, “Somatic dry-mass analysis”).

After adjusting for dry mass, somatic C:N, C:P, and N:P ratios all tended to be lower in HU guppies, although their 95% credible intervals included zero (Figure 5A). Estimated HU–LU differences were −4.2% for C:N (95% CrI [−12.4, +5.8]; *Pr*[HU < LU] = 0.898), −16.6% for C:P (95% CrI [−32.8, +3.8]; *Pr* = 0.957), and −12.8% for N:P (95% CrI [−29.8, +10.7]; *Pr* = 0.910). These trends suggest that HU guppies may have relatively more P per unit C and N in their somatic tissues, although uncertainty prevents firm conclusions about differences between urbanization origins.

**Figure 5.**
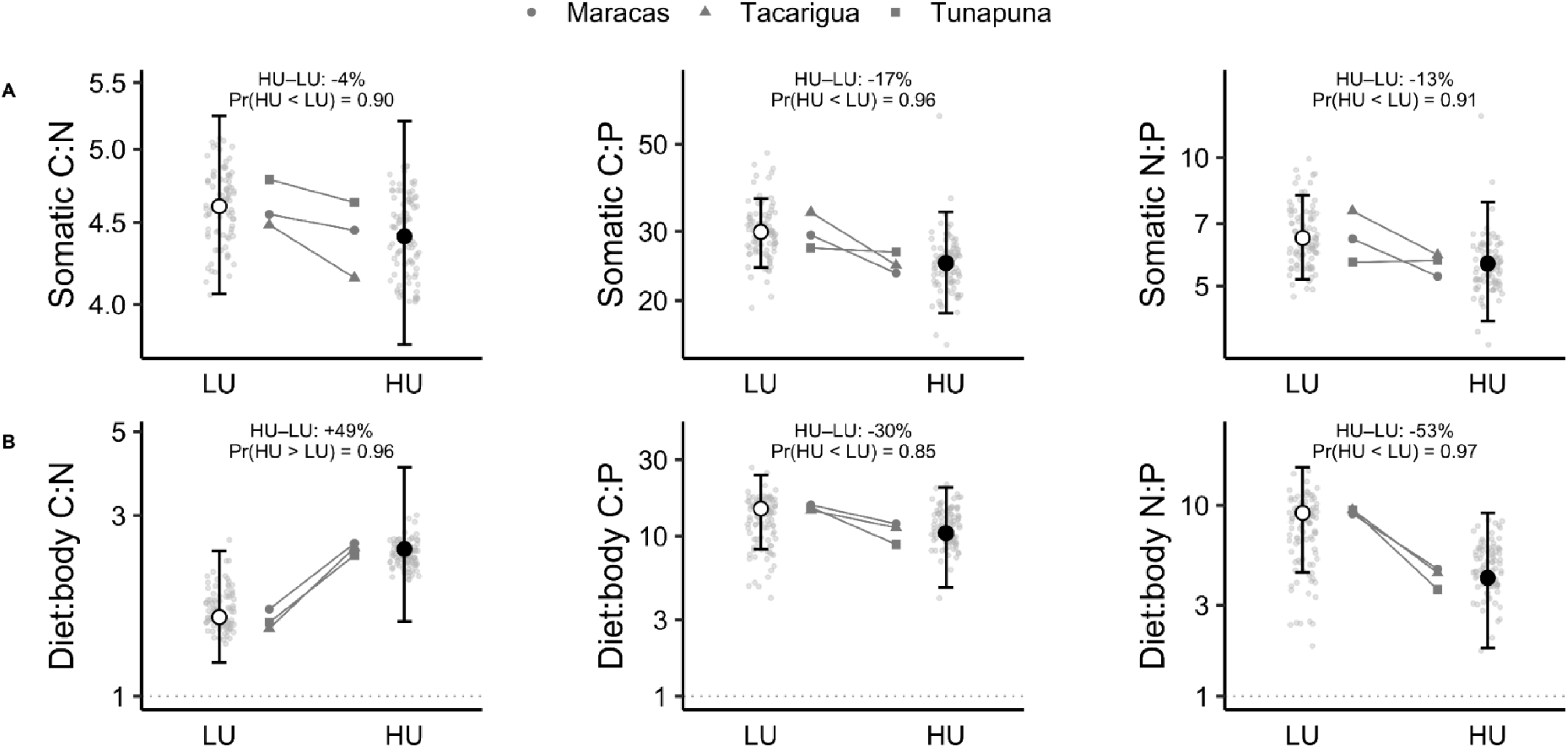
Somatic stoichiometry (A) and diet:body stoichiometric imbalance (B) of guppies from LU and HU sites. Light grey points are individual fish. Grey symbols are stream means (geometric means of the fish in each stream); shapes identify the river, and lines join the LU and HU sites of each river. White (LU) and black (HU) circles show origin-level posterior medians with 95% credible intervals. In (A), these are somatic molar ratios predicted by mixed models at the pooled mean dry mass. In (B), imbalance is the origin-level reconstructed diet ratio divided by the somatic ratio, so a value of 1 (dotted line) indicates a diet that matches body composition and values above 1 indicate a diet richer in the first element than the body. For individual fish, imbalance uses each fish’s posterior median diet ratio and its measured somatic ratio. Text in each panel gives the posterior median HU versus LU difference and the posterior probability of its direction. All y axes are on a logarithmic scale.

Dietary and somatic C:P and N:P contrasts showed similar directional trends, but somatic differences were smaller, with medians about 40% (C:P) and 20% (N:P) as large as their dietary counterparts (Figure 5). Consistent with this pattern, the diet-to-body index indicated larger declines in dietary than somatic N:P (index contrasts: −53.4%, 95% CrI [−79.8, +6.5], *Pr* = 0.965) and C:P (−29.6% [−64.9, +37.8], *Pr* = 0.855). In contrast, dietary C:N tended to increase while somatic C:N decreased, yielding a positive index contrast (+48.6% [−5.9, +145.1], *Pr* = 0.957). However, all three index contrasts had credible intervals overlapping zero.

Elemental concentrations showed broadly similar, although uncertain, patterns. Reconstructed dietary C and N concentrations tended to be lower in HU than LU guppies (−7.31 and −2.21 percentage points, respectively), whereas dietary P tended to be slightly higher (+0.044 percentage points). Somatic concentrations showed the same directional patterns, with lower C (−6.0%) and N (−1.9%) but higher P (+12.6%) in HU guppies. However, all 95% credible intervals included zero (Table S20)

## DISCUSSION

Across paired high- and low-urbanization streams, changes in guppy dietary stoichiometry were associated more strongly with shifts in assimilated resource composition than with consistent changes in resource elemental quality. Guppies from highly urbanized sites assimilated less material from aquatic invertebrates and tended to assimilate more from detrital resources, despite the absence of uniform urbanization-associated changes in resource C:N. These shifts in dietary pathways contributed particularly to differences in dietary N, but were only partly reflected in guppy somatic stoichiometry. Together, our findings provide stronger support for the Diet Composition Hypothesis than for the Resource Quality Hypothesis, particularly for dietary C:N and N:P ratios, although both mechanisms may contribute to C:P differences. Thus, the stoichiometric consequences of urbanization depend not only on the elemental composition of available resources, but also on which resources consumers assimilate and how the resulting dietary differences are reflected in consumer tissues.

### Urbanization redirects assimilated trophic pathways

The most consistent dietary difference between urbanization origins was a lower contribution of aquatic invertebrates to the assimilated diets of HU guppies. Although detrital contributions tended to increase, neither filamentous algae nor epilithon showed clear directional differences. Human modifications of streams can alter resource availability and food-web structure through changes in benthic invertebrate communities and basal resources (Price et al., 2019; Yule et al., 2015). Similar changes in resource use have been documented in omnivorous fishes inhabiting human-modified subtropical streams, where some populations increased their assimilation of terrestrial invertebrates and sedimentary organic matter (Neves et al., 2024). In our study, HU sites generally had lower canopy cover and greater watershed development than their paired LU sites, conditions that may have altered the relative availability of different food resources. Although we cannot distinguish changes in resource availability from changes in feeding behavior, both mechanisms could redirect the pathways through which guppies acquire nutrients.

The interpretation of these dietary differences also depends on assumptions about trophic discrimination, which can substantially influence estimates from stable-isotope mixing models (Bond & Diamond, 2011). Although the median invertebrate contrast remained negative under all three fixed-discrimination alternatives, its magnitude and posterior support were weaker than under the primary model, while detrital contrasts varied in direction. The relatively high carbon discrimination factor estimated by our primary model (2.82‰) may reflect physiological discrimination, differences in dietary isotopic composition, or incomplete characterization of dietary sources (Busst & Britton, 2016; Dennis et al., 2010). Consequently, although our primary model supports lower estimated invertebrate contributions in HU guppies, the magnitude of this difference remains dependent on discrimination assumptions.

### Dietary stoichiometry reflects both resource quality and resource use

Basal-resource stoichiometry did not show the uniform enrichment pattern expected under a simple nutrient-enrichment model. Instead, the elemental composition of epilithon, filamentous algae, detritus, and macroinvertebrates varied in resource-specific ways, suggesting that urbanization-associated environmental changes do not translate into a consistent improvement in food quality. Such heterogeneity is consistent with evidence that stoichiometric responses depend on resource identity and local environmental conditions (Feng et al., 2023; Sistla et al., 2015). Consequently, changes in resource stoichiometry alone may be insufficient to predict how urbanization alters the elemental composition of consumer diets.

For omnivorous consumers, however, dietary stoichiometry depends not only on the elemental composition of individual resources but also on their relative contributions to the assimilated diet. Our decomposition indicated that changes in assimilated resource mixtures contributed more strongly than changes in resource elemental quality to reconstructed dietary C:N and N:P differences. The relative importance of these mechanisms nevertheless depended on the elemental ratio considered and, particularly for P-related ratios, on how macroinvertebrate taxa were weighted. Because taxonomic turnover within broad resource categories is incorporated into our resource-quality component, these results also suggest that changes in prey composition may influence dietary stoichiometry even without changes in the relative assimilation of major resource categories.

Changes in assimilated trophic pathways did not affect dietary elemental ratios uniformly because resources differed in their relative contributions of C, N, and P (Lauridsen et al., 2014). Consequently, the relative contribution of a resource to assimilated dietary material need not match its contribution to individual elements (Higgins et al., 2006; Lauridsen et al., 2014). In HU guppies, reduced contributions of aquatic invertebrates and greater estimated contributions from detrital pathways were associated with a tendency toward lower reconstructed dietary nitrogen concentrations. Because N declined proportionally more than C, dietary C:N ratios tended to be higher in HU guppies. In contrast, dietary P showed no clear difference between urbanization origins, despite substantial changes in the trophic pathways contributing this element. Thus, lower dietary N relative to comparatively unchanged P provides a plausible explanation for the lower dietary N:P ratios in HU guppies. Although the magnitude and support for these contrasts depended on model assumptions, their differing directions underscore why changes in dietary C:N alone cannot adequately characterize the stoichiometric consequences of urbanization.

### Dietary shifts were only partly reflected in guppy body stoichiometry

Although reconstructed dietary stoichiometry differed between urbanization origins, the corresponding differences in guppy somatic tissues were smaller and more uncertain. Somatic C:N showed no clear correspondence with dietary C:N, which tended to shift in the opposite direction, although the dietary contrast was uncertain and sensitive to discrimination assumptions. Dietary and somatic N:P ratios favored the same direction, but the somatic contrast remained uncertain. Consistent with these patterns, diet-to-body indices tended to differ between urbanization origins, particularly for C:N and N:P ratios, although their credible intervals overlapped zero. Thus, although our results suggest that differences in reconstructed dietary stoichiometry were not consistently accompanied by proportional differences in somatic elemental composition, uncertainty in the diet-to-body contrasts prevents us from establishing how strongly dietary and somatic responses differed. These findings are consistent with previous observations of variable dietary–somatic correspondence in guppies (El-Sabaawi et al., 2012; Zandonà et al., 2024) and complement experimental evidence that dietary phosphorus enrichment can alter somatic elemental composition (Dalton et al., 2017).

One possible explanation is that differences in elemental assimilation, retention, and excretion modify the relationship between dietary composition and nutrient incorporation into consumer tissues (Anderson et al., 2005; Frost et al., 2005; Persson et al., 2010). These processes contribute to stoichiometric homeostasis, allowing consumers to regulate tissue elemental composition despite variation in dietary nutrient supply (López-Sepulcre et al., 2024). Differences in lipid storage could also influence somatic C:N, because lipids are carbon-rich (Czamanski et al., 2011; Downs et al., 2016; Hoffman et al., 2015). Although we did not measure these physiological processes, they offer potential explanations for why reconstructed dietary differences were not consistently reflected in guppy tissues. Importantly, our reconstructed dietary ratios do not account for differences in elemental assimilation efficiency among resources and therefore cannot establish how much of each element is ultimately incorporated into consumer biomass. Experimental comparisons of HU and LU guppies across diets differing in elemental composition, coupled with measurements of assimilation efficiency, retention, and excretion, could clarify whether urbanization is also associated with differences in stoichiometric regulation and its potential consequences for nutrient recycling.

### Implications for nutrient acquisition in changing food webs

Changes in trophic resource use may have nutritional consequences even when the dietary concentration of an individual element changes little. Under our primary reconstruction, aquatic invertebrates contributed substantially less dietary phosphorus to urban guppies, whereas detrital resources contributed more, despite no clear directional difference in overall dietary phosphorus concentrations. However, similar dietary phosphorus concentrations may not represent equivalent nutritional conditions, because bulk elemental measurements do not capture differences in the biochemical composition of food resources or the efficiency with which consumers acquire nutrients from them (Anderson et al., 2004). Thus, changes in the trophic pathways supporting omnivorous consumers may influence nutrient acquisition in ways that cannot be inferred from dietary elemental concentrations alone. In urban streams, understanding the nutritional consequences of changing resource use may therefore require considering not only the elemental composition of consumer diets, but also the resources supplying those elements and their nutritional accessibility.

## Acknowledgements

We acknowledge that the lands and waters where samples were collected for this research are the ancestral homelands of the First Peoples of Trinidad and Tobago, including Arawakan-speaking peoples and the Kalinago. We recognise the impacts of colonisation and honour the resilience and enduring presence of their descendants, particularly the Santa Rosa First Peoples Community in Arima. We are grateful to Dr. Aidan D. Farrell at the University of the West Indies, St. Augustine, for logistical support, laboratory space, and housing during field collections. We also thank Dr. Caleb Axelrod, Faith Stemmler, and Kathryn Ann Talbert for assistance in the field. We thank Dr. Eugenia Zandonà, Dr. Piatã Marques, Dr. Therese Frauendorf, Dr. Meredith Holgerson, Katherine Stenehjem, Coral del Mar Valle Rodriguez, Udita Bansal, Sylvana B. Ross, Elias Johnson, and Laurel Emanuel for valuable discussions, scientific feedback, and advice that helped shape this research. We are grateful to Kimberlee L. Sparks and Dr. Jed Sparks for their invaluable help in troubleshooting the isotope analyses and for their technical and theoretical advice on sample preparation. We are especially grateful to Dr. Rana El-Sabaawi and Dr. Alexander S. Flecker for their thoughtful guidance, constructive feedback, and valuable insights as members of J.R.A.’s dissertation committee. Funding was provided by Cornell University start-up funds to ALS and SPG and by the Cornell Atkinson Center for Sustainability’s Sustainable Biodiversity Fund Grant to JRA.

## Author Contributions

Jeferson R. Amaral, Amy Deacon, Swanne Gordon, and Andrés López-Sepulcre conceived the study and designed the methodology; Jeferson R. Amaral, Amina Mohamed, Peter C. Searle, Stephanie M. Tran, and Jillon Lewis collected the data; Jeferson R. Amaral and Andrés López-Sepulcre analyzed the data; Jeferson R. Amaral, Swanne Gordon, and Andrés López-Sepulcre led the writing of the manuscript. All authors contributed critically to the drafts and gave final approval for publication.

## Ethics Statement

All procedures complied with institutional and national guidelines for animal care and were approved by Cornell University’s Institutional Animal Care and Use Committee (IACUC protocol #2022-0170).

## Data Availability Statement

The data, R code, saved Bayesian model results, and analysis reports supporting this study are publicly available through Zenodo at https://doi.org/10.5281/zenodo.24238242. Researchers using these materials should cite the corresponding Zenodo record and the associated scientific publication when available. Researchers interested in conducting additional analyses or developing collaborative projects using these datasets are encouraged to contact the corresponding author.

## Preprint Status

This manuscript has been submitted for consideration as a Research Article in *Functional Ecology* and has not yet undergone peer review for publication in that journal.

