## Supplementary Material for "Tracing urban diet shifts into guppy stoichiometry in Trinidadian streams"

*Fish Community Analysis*

To evaluate whether HU and LU sites differed in fish community composition, we analyzed presence-absence data for all fish species recorded at each site. We calculated pairwise community dissimilarity using the Jaccard index, which is appropriate for presence-absence data. We then tested whether community composition differed between urbanization origins using a permutational multivariate analysis of variance (PERMANOVA; function "*adonis2*" from R package vegan). Because sites were paired within rivers, we constrained permutations within river identity. We also tested whether multivariate dispersion differed between urbanization origins with permutational multivariate analysis of dispersion (PERMDISP; function "*betadisper*" from R package vegan) because differences in dispersion can influence PERMANOVA results.

Fish community composition did not differ significantly between HU and LU sites based on Jaccard dissimilarity (Figure S5). Urbanization origin explained approximately 17% of the variation in community composition, but this effect was not statistically significant under the restricted permutation design ($R^2 = 0.170$, $F_{1,4} = 0.822$, $p = 0.25$). Dispersion did not differ significantly between urbanization origins ($F_{1,4} = 0.572$, permutation $p = 0.401$). Because the number of possible restricted permutations was small, the test used complete enumeration of the available permutations.

*Environmental characterization, urbanization assessment, and principal component analysis*

To characterize urbanization, we quantified environmental variables at both the watershed and local scales. At the watershed scale, we delineated watersheds and stream networks from a digital elevation model using PCRaster (Karssenberg et al., 2010). We then quantified land-cover composition (tree cover, grassland, and built-up area), impervious surface cover, and road density using ESA WorldCover data (Zanaga et al., 2022), and the Global Artificial Impervious Area dataset (Gong et al., 2020), respectively. Spatial analyses were conducted in QGIS.

At the local scale, we quantified human occupancy, canopy cover, channel structure, and water quality, including nutrient concentrations. We used raster-derived human occupancy to evaluate the consistency of our a priori urbanization classification, which followed the high- and low-urbanization threshold previously applied in Trinidadian guppy populations (Rigdon et al., 2026). Human occupancy was estimated from the LandScan Mosaic raster (Adams et al., 2025). Because occupancy estimates depend on spatial scale, we extracted values within multiple buffer radii (0.01–1 km) around each site. For each radius, we fitted a linear mixed-effects model with urbanization category (HU or LU) as a fixed effect and river, site identity, and year as random intercepts. These random effects accounted for spatial variation among river systems, repeated measurements within sites, and temporal variation in human occupancy. We compared each model against a corresponding null model excluding urbanization using likelihood-ratio tests and calculated Akaike's information criterion (AIC) for each fitted model.

We measured canopy cover using a spherical densiometer along seven transects spaced 5 m apart in a 30-m reach. At each site, we also measured water temperature (°C), pH, conductivity ($\mu S\ cm^{-1}$), and dissolved oxygen (%), total dissolved nitrogen, ammonium ($NH_4^+$), nitrate ($NO_3^-$), nitrite ($NO_2^-$), total phosphorus, phosphate ($PO_4^{3-}$), and fecal coliform abundance (*Escherichia coli*, MPN 100 $mL^{-1}$). As an index of channelization, we quantified wetted-channel area within the 30-m reach from measurements of wetted width along evenly spaced transects.

Watershed characteristics, habitat structure, and water chemistry were summarized at the site level and treated as descriptive environmental context rather than as independent inferential tests of urbanization. We additionally summarized environmental variation among sites using principal component analysis (PCA). All environmental variables were centered and scaled prior to analysis. Because the study included only six sites, PCA scores were used descriptively to characterize multivariate environmental variation among sites and were not used as predictors in the primary analyses.

Raster-derived human occupancy was consistent with our a priori classification of sites into high- and low-urbanization categories. Among the candidate buffer radii, the model using the 0.8 km buffer had the lowest AIC (375.50), followed closely by the 1.0 km buffer (AIC = 376.17). At 0.8 km, HU sites had substantially greater estimated human occupancy than LU sites (HU = 3930.77; LU = 1591.42; β = 2339.35 ± 411.90 SE, $t = 5.68$, $p = 0.030$).

Environmental contrasts between HU and LU sites were also observed in watershed development and canopy structure (Table 2). HU sites generally had lower tree and canopy cover and greater impervious and built-up cover than their paired LU sites. Differences in grassland cover, road density, dissolved nutrients, and fecal coliform abundance were less consistent among rivers.

The first three PCA axes explained 92.1% of environmental variation among sites. PC1 (59.5%) represented a nutrient-enrichment gradient, PC2 (19.1%) reflected variation in nitrogen and benthic habitat, and PC3 (13.5%) represented an open-canopy water-quality gradient. Separation between HU and LU sites was greatest along PC3, whereas differences along PC1 and PC2 were comparatively small. This suggests that differences between urbanization categories were more closely associated with local canopy openness and water-quality conditions than with the nutrient-enrichment gradient captured by PC1.

*Resource collection and processing*

To characterize epilithic algal communities, we used a BenthoTorch® fluorometer (bbe Moldaenke GmbH, Kiel, Germany) to estimate *in vivo* chlorophyll *a* concentrations (µg $cm^{-2}$) of diatoms, green algae, and cyanobacteria. We then scrubbed three submerged rocks and filtered the resulting slurry onto pre-combusted Whatman GF/F (25 mm) filters. We quantified cyanobacterial relative abundance as the proportion of total chlorophyll *a* attributed to cyanobacteria. We also sampled filamentous algae and aquatic macroinvertebrates at each site. For the filamentous algae, we carefully removed attached particulate matter and macroinvertebrates. We sampled aquatic macroinvertebrates using a 200-µm mesh Hess sampler. In the laboratory, we identified them to the lowest practical taxonomic level and dissected each individual to remove its gut before processing (Albrecht et al., 2021).

*Exploratory associations between environmental variables and resource stoichiometry*

To explore associations between environmental conditions and basal-resource stoichiometry, we fitted linear mixed-effects models (LMMs) separately for epilithon, filamentous algae, detritus, and aquatic macroinvertebrates. Molar C:N, C:P, and N:P ratios were log-transformed before analysis. For epilithon, we evaluated associations with total chlorophyll *a* concentration, the relative abundance of cyanobacteria, and environmental PCA scores. For filamentous algae and macroinvertebrates, we examined associations with environmental PCA scores. Detritus models additionally included organic matter type (CPOM, FPOM, or sewage-derived material) and its interactions with environmental gradients where applicable. River and Site were included as random intercepts in all models, with taxon identity included as an additional random intercept for macroinvertebrates. These analyses were exploratory because environmental PCA scores represented site-level conditions across only six sampling sites. Accordingly, estimated associations were interpreted descriptively rather than as independent tests of environmental effects.

Basal-resource stoichiometry exhibited resource-specific associations with environmental gradients (Table S1; Figure S4). Epilithon C:N ratios increased along PC3, the open-canopy water-quality gradient, but decreased with increasing total chlorophyll concentration. Higher cyanobacterial relative abundance also tended to be associated with lower epilithon C:P and N:P ratios. Filamentous algae showed contrasting associations with PC2 and PC3, with C:P ratios increasing along PC2 but decreasing along PC3. Filamentous algal C:N ratios also decreased along PC3, whereas associations involving N:P ratios were less pronounced. However, none of the filamentous algal associations was statistically supported (all $p > 0.18$), and these models had few approximate degrees of freedom (about 2 to 3). Macroinvertebrate stoichiometry showed comparatively limited variation along these gradients, with C:N ratios decreasing along PC3 but no clear associations for C:P or N:P ratios.

Detrital stoichiometry varied among organic matter types and environmental gradients. CPOM C:N ratios tended to decrease along PC1 ($p = 0.062$), whereas FPOM C:N and C:P ratios decreased along PC3. In contrast, sewage-derived material tended to show a positive association between C:N ratios and PC3 ($p = 0.072$). Compared with CPOM, sewage-derived material also had substantially lower C:P and N:P ratios, and C:P and N:P ratios of all detrital types decreased along PC1. Overall, these exploratory analyses indicated that associations between environmental conditions and resource stoichiometry differed among resource types, elemental ratios, and organic matter pools, rather than following a consistent direction across the food web.

*Somatic dry-mass analysis*

To assess whether somatic dry mass differed between urbanization origins, we fitted a linear mixed-effects model with log-transformed somatic dry mass as the response, Origin as a fixed effect, and River and Site as random intercepts. HU guppies tended to have lower somatic dry mass (26.4 ± 17.4 mg, mean ± SD) than LU guppies (48.0 ± 31.8 mg; $\beta = -0.520 \pm 0.192$ SE on the log scale, $t_2 = -2.71$, $p = 0.11$). Geometric mean dry mass was lower at the HU site in all three rivers (Maracas: −17%; Tacarigua: −40%; Tunapuna: −57%). Because the design was balanced, this model is equivalent to a paired t-test of site mean log dry mass across the three rivers, with two degrees of freedom.

*Hierarchical Bayesian stable isotope mixing models*

We estimated the assimilated dietary composition of Trinidadian guppies using hierarchical Bayesian stable isotope mixing models implemented in JAGS. The analyses included 180 guppies collected from six sites, with one HU and one LU site in each of three rivers. We considered four broad dietary sources: detritus, epilithon, filamentous algae, and aquatic invertebrates. Rather than estimating dietary composition independently for each site, we used a hierarchical framework that allowed resource isotope signatures and consumer diets to vary among rivers and sites through partial pooling, while estimating resource isotope means and variances jointly with dietary proportions (Semmens et al., 2009; Ward et al., 2010).

*Modeling resource isotope signatures*

Resource isotopes were estimated using 292 analytical records: 47 detritus, 18 epilithon, 18 filamentous algae, and 209 invertebrate records. Each record was retained as an individual observation in the source likelihood, rather than reducing the data to predetermined site-specific means and standard errors. Because some analytical records represent pooled biological material, the modeled variation describes differences among analytical records and does not necessarily represent variation among individual organisms.

We modeled the mean $\delta^{13}C$ and $\delta^{15}N$ of each broad dietary source using a hierarchical structure consisting of an overall source mean, river-specific deviations, and site-specific deviations. This partial-pooling approach allowed resource isotope signatures to vary spatially while incorporating information from the broader dataset. Source isotope variability was also estimated hierarchically, with source- and isotope-specific variation and partially pooled site-level standard deviations.

To account for heterogeneity within dietary sources, we retained the identities of individual invertebrate taxa and detritus components. The dataset contained 26 distinct invertebrate taxa and three detritus categories (CPOM, FPOM, and sewage-derived organic matter), representing 62 and 16 site-specific subgroup combinations, respectively. The model estimated isotope deviations among these subgroups while maintaining a single broad dietary contribution for invertebrates and a single contribution for detritus. Because epilithon and filamentous algae were not subdivided into finer resource categories, additional between-subgroup deviations were not estimated for these sources, although their site-specific isotope means and analytical variation remained explicitly modeled.

Within each site and broad source, subgroup isotope deviations were centered using weights proportional to the number of analytical records available for each subgroup. This centering preserved the analytical-record-weighted isotope centroid of each broad source within a site while allowing variation among its constituent subgroups. Importantly, these weights represented the distribution of analytical measurements rather than resource abundance, availability, or taxon-specific consumption. Consequently, the model estimated assimilated contributions from the four broad source categories, not the consumption of individual invertebrate taxa or detritus components. These weights defined the centering of resource subgroups within the source likelihood. They did not represent dietary contributions or determine the gut-informed prior, although their influence on estimated source signatures could indirectly affect dietary inference.

*Modeling consumer dietary composition*

Consumer dietary proportions were modeled using an isometric log-ratio (ILR) transformation, which permitted the four source contributions to vary while remaining positive and summing to one. We used a sequential orthonormal ILR basis contrasting detritus with epilithon, their combined composition with filamentous algae, and the first three sources with invertebrates. The dietary model included urbanization origin (HU or LU) as a fixed effect, river and site as random effects, and standardized log-transformed somatic dry mass as a covariate.

Fish $\delta^{13}C$ values were adjusted for lipid content using mathematical normalization (Post et al., 2007). We also centered $\delta^{15}N$ measurements of both consumers and resources relative to the mean snail $\delta^{15}N$ measured at each site. Expected consumer isotope signatures were calculated from the estimated dietary proportions and site-specific source isotope signatures, accounting for measured source C and N concentrations and trophic discrimination factors (TDFs). Site-specific source elemental concentrations and snail-derived nitrogen baselines were treated as fixed inputs. Consumer isotope observations were modeled jointly using a multivariate normal distribution, allowing correlated C and N residuals and partially pooled site-specific residual variability.

The resource and consumer likelihoods were fitted jointly. Accordingly, resource isotope observations informed estimates of source signatures, while consumer isotope observations could also update those estimates through the dietary mixing model. Because four dietary sources were estimated using two isotopes, dietary contributions were not uniquely determined by isotopic signatures alone; the posterior estimates also depended on the source model, concentration weighting, dietary priors, and discrimination assumptions.

We specified a single gut-content-informed Dirichlet prior for the global baseline dietary composition, with concentration parameters of 2.033, 1.068, 1.652, and 1.247 for detritus, epilithon, filamentous algae, and invertebrates, respectively (total concentration = 6). To construct this prior, food-item occurrence frequencies were summarized within available river-by-origin groups, averaged within each origin, and then averaged equally across LU and HU populations, with a small positive occurrence weight assigned to epilithon (0.05). The resulting frequencies were rescaled to sum to one, and each concentration parameter was set to 1 plus twice the corresponding rescaled frequency. This procedure produced a single baseline prior rather than separate origin-specific dietary priors. Baseline proportions were transformed into ILR coordinates, with LU as the reference category and HU represented by an additional zero-centered normal effect on each ILR coordinate. Thus, no directional urbanization effect was imposed in ILR space, although the reference-category parameterization induced different marginal prior distributions of LU and HU dietary proportions. Because gut-content observations were unavailable for Tunapuna LU, the prior was constructed from the available river-by-origin groups.

To assess whether dietary estimates depended on the gut-content-informed baseline prior, we fitted an alternative model using a uniform Dirichlet(1, 1, 1, 1) prior. This specification assigned equal prior concentration parameters to the four dietary sources, without incorporating information from gut-content observations. All other model components were unchanged. We compared posterior dietary contributions, equal-weighted within-river HU−LU contrasts, and directional posterior probabilities between the two prior specifications.

To evaluate sensitivity to this reference-category parameterization, we fitted an alternative model using symmetric urbanization coding. In the original specification, the Dirichlet-derived ILR coordinates represented the LU baseline, with the urbanization coefficient added for HU populations. In the alternative specification, the same global baseline was treated as the midpoint between origins, with $-\beta/2$ applied to LU and $+\beta/2$ to HU for each ILR coordinate. Both specifications retained the same zero-centered Normal(0, 1) prior on the HU–LU coefficient, and all other likelihood components, data inputs, and prior distributions were unchanged. This comparison evaluates whether differences in the induced marginal dietary priors affected the estimated urbanization contrasts.

We additionally evaluated the influence of the prior scale assigned to site-level dietary random effects. The primary model assigned the dietary site-effect standard deviation a half-normal prior with scale 0.35 on the ILR scale. In an alternative specification, we doubled this scale to 0.70, allowing greater variation in dietary composition among sites. All other model components, including the source hierarchy, gut-content-informed dietary prior, LU-reference urbanization coding, and jointly estimated trophic discrimination factors, were retained. We compared posterior dietary contributions, equal-weighted within-river HU−LU contrasts, estimated site- and river-level dietary standard deviations, and model diagnostics between the two specifications.

The dietary, resource, and residual variance parameters were assigned regularizing prior distributions, with their scales defined on the ILR, standardized isotope, or log-standard-deviation scales as appropriate. Complete prior specifications are provided in Table S2.

*<u>Prior specification and sensitivity of resource elemental composition models</u>*

We used empirically calibrated, weakly informative priors for the Bayesian hierarchical models estimating carbon, nitrogen, and phosphorus concentrations in epilithon, filamentous algae, invertebrates, and detritus. Because elemental concentrations differed substantially among resources, intercept prior centers were calculated separately for each resource-element combination using the equally weighted mean of the six site-level averages on the transformed scale. We assigned normal intercept priors with a standard deviation of 0.85 for log-Gaussian models and 1 for the beta model of detritus carbon. The urbanization effect was assigned a zero-centered normal prior (mean = 0, SD = 0.7), independently of the observed HU–LU contrast. Random-effects standard deviations were assigned half-normal priors with scales of 0.35 for River and Site and 0.5 for invertebrate Taxa. For log-Gaussian models, residual standard deviations were assigned half-Student-t priors with three degrees of freedom and scales calibrated to 1.5 times the median within-site standard deviation, bounded between 0.25 and 1.5. The beta model of detrital carbon was assigned a Gamma(2, 0.1) prior (shape, rate) on its precision parameter.

We evaluated the sensitivity of resource elemental HU–LU contrasts to alternative intercept prior centers and wider priors for the urbanization effect and variance components. Alternative intercept centers were shifted by ±log(2) on the transformed scale, whereas prior scales for selected fixed effects and variance components were doubled individually. We also compared models with and without Site as an additional random intercept. Posterior contrasts were generally insensitive to changes in prior specification, although some estimates, particularly detrital phosphorus, exhibited greater sensitivity. Including Site increased the widths of 95% credible intervals by 1.34–2.92 times while leaving median contrasts largely unchanged. Thus, accounting for site-level clustering had a greater influence on estimated uncertainty than the alternative prior specifications examined. We consequently retained the site-adjusted hierarchical models with empirically calibrated priors for subsequent dietary reconstruction.

*Trophic discrimination estimation and sensitivity analyses*

Trophic discrimination factors (TDFs) introduce an important source of uncertainty into isotope-based dietary estimates because they can vary with consumer identity, tissue composition, and diet. Experimental studies have demonstrated diet-dependent discrimination in guppies and other freshwater fishes, while recent meta-analyses indicate substantial variation among taxa, tissues, trophic levels, and dietary sources (Busst & Britton, 2016; Dennis et al., 2010; Stephens et al., 2023). Given this variability and the absence of an experimental TDF calibration for the guppy somatic tissues analyzed here, we used a primary model in which carbon and nitrogen discrimination factors were estimated jointly with source isotope signatures and dietary proportions.

The primary model included two globally shared TDF parameters with independent normal priors informed by literature-based estimates (carbon: 0.40 ± 1.30‰; nitrogen: 2.54 ± 1.27‰, mean ± SD) (Post, 2002; Vanderklift & Ponsard, 2003). These values have also been adopted in stable-isotope mixing analyses of guppies inhabiting polluted freshwater environments (de Carvalho et al., 2019). The priors represented uncertainty about the common discrimination parameters rather than experimentally calibrated variation among individual guppies. Consequently, the estimated TDFs were shared across individuals, sites, and dietary sources.

To evaluate the dependence of dietary estimates on discrimination assumptions, we fitted three additional specifications using the same hierarchical structure and all non-TDF inputs. The first used fixed TDFs of 0.40‰ for carbon and 2.54‰ for nitrogen (Post, 2002; Vanderklift & Ponsard, 2003). The remaining scenarios used estimates for omnivorous fishes from Stephens et al. (2023): a C3-derived dietary-origin scenario (carbon: 1.20‰; nitrogen: 4.10‰) and a mixed-origin scenario (carbon: 1.40‰; nitrogen: 3.10‰). These scenarios represented alternative literature-supported assumptions rather than independent calibrations for the sampled guppies.

Across the four specifications, we compared posterior estimates of absolute dietary contributions and HU–LU differences. Urbanization contrasts were calculated separately within each river and then averaged with equal weighting across the three paired rivers. These contrasts were based on site-level dietary predictions at a common reference somatic dry mass (29.68 mg, the pooled geometric mean). The resulting estimates describe the sampled river pairs conditional on their fitted site effects, rather than predictions for previously unsampled rivers. We summarized posterior distributions using medians, 95% credible intervals, and posterior probabilities of directional differences. We also examined posterior correlations between discrimination factors and absolute dietary contributions at the site and urbanization-origin levels, as well as paired HU−LU dietary contrasts, to evaluate their dependence within the primary model.

For paired dietary contrasts, we also calculated least-squares regression slopes across posterior draws to express the linear association with each TDF in percentage points per ‰. These slopes characterize within-model posterior dependence rather than the effects of fixing TDFs at alternative values.

<u>*Source-only diagnostic and model evaluation*</u>

To evaluate the influence of consumer isotope observations on the estimation of resource signatures, we fitted a source-only hierarchical model that retained the original resource observations, hierarchical source structure, subgroup definitions, weighting scheme, and source priors from the jointly estimated-TDF model, but excluded the consumer isotope likelihood and all dietary and discrimination parameters. We compared posterior source isotope means from the source-only model with those estimated under each mixing-model specification. For each of the 48 site–source–isotope combinations, we calculated the difference between posterior medians, summarized absolute deviations, and identified joint-model posterior medians falling outside the corresponding source-only 95% credible intervals. We additionally compared fitted source isotope means with observed analytical-record means to examine discrepancies between estimated and measured resource signatures. These comparisons were interpreted as diagnostics of consumer-to-source feedback and source-model behavior rather than independent validation of source estimates or TDF assumptions.

We used the jointly estimated-TDF specification as the primary analysis because it allowed literature-informed uncertainty in discrimination factors to be represented in the absence of an experimental calibration for the sampled guppy tissues. We evaluated its behavior through comparisons with the source-only model and three fixed-TDF alternatives. These analyses assessed the sensitivity of dietary estimates to discrimination assumptions and the extent of consumer-to-source feedback, rather than providing independent validation of the estimated TDFs.

All seven completed mixing-model specifications were fitted using four Markov chains of 1,000,000 iterations each, including 500,000 burn-in iterations and thinning by 100. Convergence was assessed using rank-normalized $\hat{R}$, bulk and tail effective sample sizes (ESS), and posterior predictive checks comparing observed and replicated consumer and resource isotope root mean square errors (RMSEs). Consumer checks were calculated separately for each site and isotope, and resource checks were calculated for each site, source, and isotope. Posterior predictive tail probabilities below 0.025 or above 0.975 were flagged for examination. We examined posterior associations among discrimination factors, dietary contributions, and fitted source isotope signatures to characterize potential parameter dependence. We also calculated pairwise Pearson correlations among posterior dietary source contributions at the origin level and among paired HU−LU contrasts to characterize posterior dependence among dietary estimates. These diagnostics were used to evaluate computational convergence, model adequacy, and sensitivity to modeling assumptions, rather than to establish unique identification of dietary proportions.

*Posterior dietary estimates and source correlations*

Under the jointly estimated-TDF model, posterior dietary estimates indicated differences in assimilated dietary composition between LU and HU guppies (Figure S1; Table S3). The clearest contrast involved aquatic invertebrates, which accounted for a greater proportion of the estimated diet in LU populations (posterior median: 41.3%) than in HU populations (5.6%). The equal-weighted HU−LU contrast across the three rivers was −33.7 percentage points (95% credible interval, CrI [−60.1, −5.0]), with a posterior probability of 0.991 that invertebrate contributions were lower in HU populations. Detritus exhibited the opposite pattern, with estimates suggesting greater dietary contributions in HU (54.2%) than in LU populations (19.3%). The corresponding HU−LU contrast was +33.0 percentage points (95% CrI [−4.6, 64.2]), with a posterior probability of 0.954 that detrital contributions were higher in HU populations. Across the four TDF specifications, the median HU−LU contrast for invertebrate contributions ranged from −2.6 to −33.7 percentage points, whereas the direction of the detrital contrast varied among specifications (Table S5). In contrast, epilithon and filamentous algae showed no clear directional differences between urbanization origins (Table S3). Posterior dietary contributions exhibited moderate correlations, with the strongest negative association between detritus and epilithon in HU populations ($r = -0.68$). Among paired HU−LU contrasts, correlations ranged from −0.55 to −0.15 (Figure S3), reflecting posterior dependence in the allocation of dietary contributions among sources. These relationships indicated site-dependent trade-offs between the estimated contributions of these resources, although negative correlations are also expected among compositional dietary proportions.

The reduction in estimated invertebrate assimilation was consistent in direction across all three rivers (Table S4). Posterior median HU−LU contrasts were −30.4 percentage points in Maracas, −44.1 in Tacarigua, and −24.8 in Tunapuna. The 95% credible intervals excluded zero in Maracas and Tacarigua but overlapped zero in Tunapuna. Detrital contributions were also estimated to be greater in HU populations in all three rivers, although their river-specific credible intervals included zero. These results indicate a broadly consistent directional pattern of reduced invertebrate assimilation between urbanization categories, accompanied by less certain increases in detrital contributions.

*Sensitivity to trophic discrimination factors*

The magnitude and posterior support of urbanization-associated dietary differences depended on the TDF specification (Table S5). All four models produced negative posterior median HU−LU contrasts for invertebrate assimilation, but the estimated magnitude varied considerably. The three fixed-TDF specifications yielded contrasts of −2.6 percentage points under conventional discrimination values, −4.1 under the Stephens C3-derived scenario, and −14.5 under the Stephens mixed-origin scenario, compared with −33.7 under the jointly estimated-TDF model. Corresponding posterior probabilities of lower invertebrate assimilation in HU populations ranged from 0.789 to 0.961 across the fixed-TDF scenarios, compared with 0.991 under the primary model. Unlike the primary model, all three fixed-TDF specifications had 95% credible intervals that included zero.

Detrital contributions were more sensitive to discrimination assumptions. Whereas the jointly estimated-TDF model supported greater detrital assimilation in HU populations, the conventional and Stephens C3 scenarios produced slightly negative median HU−LU contrasts, and the Stephens mixed-origin scenario produced a smaller positive contrast (Table S5). Thus, although the direction of the median invertebrate contrast was consistent across all four specifications, the magnitude and certainty of this contrast, as well as the inferred contribution of detritus, remained dependent on the assumed discrimination factors.

*Sensitivity to urbanization-prior parameterization*

Dietary contrasts were relatively stable under symmetric urbanization coding (Table S12). The median HU–LU difference in invertebrate contributions changed from –33.7 percentage points (95% CrI [−60.1, −5.0]) under the original LU-reference specification to –38.9 percentage points [−65.0, −7.2] under symmetric coding. The posterior probability of lower invertebrate contributions in HU populations remained 0.991 in both models. The median detrital contrast increased from +33.0 to +37.2 percentage points, and its 95% CrI [+1.3, +67.3] no longer overlapped zero. Differences between specifications were smaller for epilithon and filamentous algae, with changes in their median contrasts of less than three percentage points. Thus, the direction and posterior support of the invertebrate contrast were relatively insensitive to the origin-prior parameterization examined here, although its magnitude remained dependent on TDF assumptions.

The posterior median dietary site SD was similar under LU-reference (0.487, 95% CrI [0.034, 1.053]) and symmetric-origin coding (0.494, 95% CrI [0.040, 1.02]), suggesting that the alternative parameterization had little effect on estimated site-level dietary heterogeneity.

<u>*Sensitivity to dietary prior specifications*</u>

Widening the dietary site-effect prior produced broadly similar urbanization contrasts (Table S13), although the posterior estimates of spatial heterogeneity changed substantially. The median HU−LU contrast for invertebrate contributions changed from −33.7 percentage points (95% CrI [−60.1, −5.0]; Pr[HU < LU] = 0.991) under the primary prior to −40.1 percentage points ([−67.4, −7.1]; *Pr*[HU < LU] = 0.992) under the wider prior. The corresponding detrital contrast changed only slightly, from +33.0 [−4.6, 64.2] to +33.6 percentage points [−4.2, 68.0]. Epilithon and filamentous algae continued to show uncertain directional differences between origins.

The posterior median invertebrate contrast remained negative in all three rivers under the wider prior: −38.2 percentage points in Maracas (95% CrI [−87.3, −0.4]), −58.1 in Tacarigua [−92.1, −1.7], and −24.7 in Tunapuna [−59.9, 16.2]. The posterior median dietary site-effect SD increased from 0.487 (95% CrI [0.034, 1.053]) to 1.015 [0.199, 1.968], whereas the dietary river-effect SD decreased from 0.512 [0.031, 1.227] to 0.362 [0.017, 1.103]. Estimated discrimination factors were similar between specifications, with posterior medians of 2.82‰ and 2.86‰ for carbon and 3.00‰ and 2.99‰ for nitrogen under the primary and wider priors, respectively. The wider-prior model exhibited satisfactory convergence (maximum core $\hat{R}$ = 1.003; minimum bulk ESS = 977) and no extreme posterior predictive RMSE probabilities among the 12 consumer or 48 resource comparisons. Thus, the overall invertebrate contrast was relatively stable across the two site-effect priors, despite considerable sensitivity in the inferred magnitude of site-level dietary heterogeneity.

Replacing the gut-informed Dirichlet prior with a uniform Dirichlet(1, 1, 1, 1) prior produced nearly identical HU−LU contrasts for invertebrates (−33.7 versus −34.0 percentage points) and detritus (+33.0 versus +32.6 percentage points; Table S14).

*Trophic discrimination estimates and source-model diagnostics*

The jointly estimated model yielded posterior median discrimination factors of 2.82‰ for carbon (95% CrI [2.20, 3.33]) and 3.00‰ for nitrogen ([2.72, 3.41]; Table S7). The carbon estimate was substantially higher than its literature-informed prior mean of 0.40‰, whereas the nitrogen estimate remained closer to its prior mean of 2.54‰. Although elevated relative to conventional carbon discrimination values, the estimated carbon TDF falls within experimentally documented ranges for omnivorous freshwater fishes (Busst & Britton, 2016). Experimental studies of guppies have also demonstrated that carbon discrimination varies with dietary isotope composition (Dennis et al., 2010), supporting the biological relevance of accounting for discrimination uncertainty, although neither study independently calibrates the TDF estimated here.

Posterior correlations between discrimination factors and paired HU−LU dietary contrasts were modest (Pearson $|r| \leq 0.224$; Table S8), although regression slopes indicated associations with contrast magnitude. For invertebrates, the HU−LU contrast was negatively associated with carbon discrimination (−8.10 percentage points per ‰) and positively associated with nitrogen discrimination (+17.76 percentage points per ‰), whereas detritus showed the opposite relationships (+11.93 and −16.36 percentage points per ‰, respectively). Associations with absolute dietary contributions were stronger at some sites, particularly between nitrogen discrimination and filamentous-algae contributions at Tunapuna LU ($r = 0.400$). At the urbanization-origin level, the strongest association also involved nitrogen discrimination and filamentous-algae contributions in LU populations (r = 0.339). Tail effective sample sizes ranged from 7,490 to 15,387 across the TDF parameters and dietary contrasts. These results indicated satisfactory posterior sampling, while the dietary estimates remained dependent on discrimination assumptions. These regression slopes describe local linear associations within the fitted posterior and should not be extrapolated to predict differences among the fixed-TDF specifications, which allow resource signatures and other model parameters to change.

Source isotope estimates generally changed little after incorporating consumer observations, although the magnitude of these changes varied among resources. Across the 48 site–source–isotope combinations, the median absolute difference between source-only and primary-model posterior medians was 0.089‰ (Table S9). The smallest shifts occurred for invertebrates, whereas filamentous algae exhibited the largest differences, reaching 2.12‰. All 48 primary-model posterior medians remained within the corresponding source-only 95% credible intervals. Compared with the fixed-TDF specifications, the primary model produced smaller mean absolute and maximum departures from the source-only estimates (Table S10). These comparisons characterize the influence of consumer observations on source estimation rather than independently validating the estimated discrimination factors. Site-specific isotope signatures of guppies and dietary resources are shown in Figure S2.

For example, at Tunapuna LU, the posterior estimate of filamentous-algae $\delta^{13}C$ was −29.80‰ in the source-only model and −28.05‰ in the primary mixing model, compared with −22.07‰ under the conventional fixed-TDF specification. This illustrates how discrimination assumptions influenced the estimated source signatures, with the primary model producing substantially less displacement from the source-only estimate in this case.

All seven mixing-model specifications exhibited satisfactory convergence, with maximum core $\hat{R}$ values ranging from 1.002 to 1.005 and minimum bulk effective sample sizes ranging from 847 to 2,297 (Table S11). Posterior predictive checks identified no extreme RMSE tail probabilities among the 12 consumer or 48 resource site-by-isotope comparisons for any specification. The source-only model also exhibited satisfactory convergence (maximum $\hat{R}$ = 1.008; minimum bulk ESS = 920).

*Sensitivity of dietary reconstruction to resource elemental prediction scale*

Most resource elemental models used a Gaussian likelihood for log-transformed concentrations. To evaluate whether back-transformation influenced reconstructed dietary stoichiometry, we compared conditional arithmetic means, calculated as $\exp(\eta + \sigma^2/2)$, with conditional medians, calculated as $\exp(\eta)$, where $\eta$ is the predicted linear response and $\sigma$ is the residual standard deviation. For detrital %C, modeled using a beta likelihood, the inverse-logit transformation provided the fitted mean under both approaches. We retained conditional arithmetic means for the primary reconstruction because dietary elemental supply was calculated by weighting and summing the elemental concentrations of assimilated sources. Both reconstructions used the same 10,000 posterior draws of site-specific dietary proportions from the hierarchical mixing model and the same resource posterior-draw pairings.

Using conditional medians produced a higher median HU–LU C:N contrast (+50.7%, 95% CrI [−6.4, +150.8]) than the primary arithmetic-mean reconstruction (+42.7% [−9.0, +132.6]). C:P contrasts remained similar (−42.2% versus −41.3%), as did N:P contrasts (−62.2% versus −59.2%). Diet composition remained the most strongly supported contributor to C:N differences (posterior probabilities of dominance: 0.954 versus 0.935) and N:P differences (0.924 versus 0.908). In contrast, neither component was consistently dominant for C:P (0.499 versus 0.472; Table S15).

*Sensitivity to detrital subtype weighting*

Because the relative contributions of CPOM, FPOM, and sewage-derived organic material to assimilated detritus were unknown, we evaluated three weighting schemes (Table S16). The primary reconstruction assigned equal weights to all three subtypes. Alternatively, site-occurrence weighting assigned equal weights to the subtypes recorded at each sampling site and averaged the resulting mixtures equally across rivers within each origin. A third scenario excluded sewage-derived material and assigned equal weights to CPOM and FPOM. All scenarios retained the same hierarchical dietary posterior, resource elemental posterior draws, and conditional arithmetic mean predictions, varying only how detrital elemental composition was reconstructed. Subtype occurrence reflected analytical sampling rather than measured environmental availability.

Site-occurrence weighting produced dietary contrasts similar to the primary reconstruction. C:N remained higher in HU diets (+41.5% versus +42.7%), whereas C:P (−42.4% versus −41.3%) and N:P (−59.6% versus −59.2%) remained lower. Excluding sewage had a greater influence on P-related ratios. The C:P contrast decreased in magnitude to −31.0% (95% CrI [−64.3, +30.4]), whereas the N:P contrast changed to −51.3% [−77.6, +1.7]. Although the latter credible interval included zero, the posterior probability of a lower N:P ratio in HU remained 0.972. Diet composition remained the more strongly supported contributor to C:N differences across weighting schemes (posterior probabilities of dominance: 0.924–0.935), whereas support was more variable for N:P (0.843–0.908) and C:P (0.378–0.472; Table S16).

*Sensitivity to hypothetical sewage enrichment*

Because the contribution of sewage-derived material to detritus was not quantified, we further evaluated hypothetical scenarios representing greater sewage contributions in HU than LU populations. We considered three hypothetical scenarios in which sewage represented 50% of detritus in HU versus 10% in LU, 70% versus 5%, and 70% versus 0%. The remaining contributions were distributed between CPOM and FPOM according to their relative site-occurrence weights. These scenarios represented assumptions about detrital elemental composition rather than measured estimates of detrital availability or assimilation. All other reconstruction inputs and posterior-draw pairings remained unchanged.

Across these scenarios, C:N ratios remained higher in HU diets (+39.0% to +39.7%), although their 95% credible intervals included zero. N:P ratios remained lower (−65.2% to −70.7%), with credible intervals excluding zero in all scenarios. C:P differences also became more negative (−50.8% to −58.6%), with credible intervals excluding zero under each hypothetical scenario. The resource-quality contribution became increasingly negative for C:P (−37.9% to −45.9%) and N:P (−30.8% to −39.1%). Consequently, the posterior probability that diet composition contributed more than resource quality declined to 0.152–0.232 for C:P and 0.712–0.796 for N:P. In contrast, diet composition remained the more strongly supported contributor to C:N differences, with probabilities exceeding 0.92 in all scenarios (Table S17).

*Sensitivity to macroinvertebrate taxon weighting*

Because macroinvertebrates included multiple taxa whose relative abundances were not quantified, we evaluated whether alternative taxonomic weighting affected reconstructed dietary stoichiometry. The primary reconstruction assigned equal weights to all 26 modeled taxa. As an alternative, we assigned equal weights to taxa recorded at each sampling site and averaged these site-specific mixtures equally across the three rivers within each origin. Taxa not recorded at a site were treated as unrepresented in its sampled mixture, without assuming ecological absence. We recalculated macroinvertebrate %C, %N, and %P concentrations from the existing taxon-specific posterior predictions, while retaining fitted taxon effects and excluding River- and Site-specific deviations. All other resource predictions, hierarchical dietary proportions, and posterior-draw pairings were unchanged.

Alternative taxonomic weighting produced relatively small changes in total dietary contrasts. C:N remained higher in HU diets (+43.2% versus +42.7% in the primary reconstruction), whereas C:P (−37.7% versus −41.3%) and N:P (−56.8% versus −59.2%) remained lower. However, the decomposition of P-related differences was more sensitive. For C:P, the resource-quality contribution became more negative under site-occurrence weighting (−28.8%, 95% CrI [−48.4, −0.2]), compared with −24.9% [−46.4, +5.3] under equal weighting. The probability that diet composition contributed more than resource quality decreased from 0.472 to 0.296. For N:P, the diet-composition contribution remained negative (−43.9%), although its 95% credible interval included zero [−72.5, +2.7], and the probability of composition dominance decreased from 0.908 to 0.826. In contrast, support for diet-composition dominance in C:N differences remained nearly unchanged (0.935 versus 0.936; Table S18).

<u>*Overall sensitivity of reconstructed dietary stoichiometry*</u>

Reconstructed dietary stoichiometry also depended on the TDFs. Under the three fixed literature TDF values, C:P and N:P ratios remained lower in HU guppies but with 95% credible intervals including zero, whereas the C:N contrast remained positive but smaller under the Stephens mixed-origin values and became slightly negative under the conventional and Stephens C3 values (Table S6). Across alternative resource-weighting assumptions, sensitivity analyses generally preserved the direction of reconstructed dietary contrasts. Diet composition remained the larger contributor to C:N differences across all resource-weighting scenarios, although not under the conventional or Stephens C3 TDF values (Table S6), whereas its relative importance for N:P weakened under alternative resource-weighting assumptions. The N:P contrast consistently favored lower values in HU guppies, although its 95% credible interval included zero when sewage-derived material was excluded from detrital elemental predictions. C:P differences were more sensitive to resource composition. Under hypothetical sewage-enrichment scenarios, the C:P contrast strengthened (all 95% credible intervals excluded zero) and was driven mainly by resource quality rather than diet composition (Supplementary Material, "Sensitivity to hypothetical sewage enrichment"; Table S17).

*Final resource elemental models and posterior predictions*

Following the prior sensitivity analyses described above, we selected the site-adjusted hierarchical models with empirically calibrated priors for estimating resource elemental composition. We fitted 12 Bayesian models, separately estimating %C, %N, and %P for epilithon, filamentous algae, aquatic macroinvertebrates, and detritus. Each model included urbanization origin as a fixed effect and river and site as random intercepts. Invertebrate models additionally included taxon identity as a random intercept, whereas detritus models included organic matter subtype as a fixed effect to distinguish coarse particulate organic matter (CPOM), fine particulate organic matter (FPOM), and sewage-derived material. Elemental concentrations were modeled using Gaussian likelihoods for log-transformed responses, except detrital %C, which was modeled using beta regression.

We fitted models using four Markov chain Monte Carlo (MCMC) chains, initially with 6,000 iterations per chain, including 3,000 warmup iterations. Models that failed convergence or sampling diagnostics were refitted using longer chains and higher target acceptance probabilities, with up to 10,000 iterations per chain. We assessed convergence using rank-normalized $\hat{R}$ and bulk and tail effective sample sizes (ESS), and examined divergent transitions and maximum tree-depth warnings. Acceptance criteria required $\hat{R} \leq 1.01$ and effective sample sizes $\geq 400$, and no maximum tree-depth warnings or divergent transitions, except for an allowance of up to two divergent transitions in the invertebrate carbon model when all other criteria were satisfied. We additionally compared posterior predictive distributions with observed elemental concentrations using 200 posterior predictive draws per model.

For each fitted model, we randomly retained 10,000 posterior draws to propagate uncertainty in resource elemental composition into dietary reconstruction. Predictions were generated separately for HU and LU origins, with River- and Site-specific random-effect deviations set to zero. For invertebrates, predictions retained the estimated taxon-specific deviations and were averaged equally across the 26 modeled taxa. Detrital predictions retained the fixed effects of organic matter subtype and were combined using equal weights for CPOM, FPOM, and sewage-derived material in the primary reconstruction. These origin-standardized elemental predictions were subsequently combined with site-specific dietary proportions from the hierarchical stable-isotope mixing model to reconstruct dietary stoichiometry and estimate paired HU–LU contrasts within each river. The 10,000 dietary draws were sampled without replacement from the 20,000 retained mixing-model draws using a fixed random seed (123), preserving posterior dependence among the four dietary source contributions within each draw.

We used conditional arithmetic mean elemental concentrations on the original percentage scale for the primary reconstruction, accounting for residual variance when back-transforming log-Gaussian predictions. Posterior estimates of %C, %N, and %P were retained separately and used to calculate molar carbon-to-nitrogen, carbon-to-phosphorus, and nitrogen-to-phosphorus ratios using the corresponding atomic masses. Because elemental concentrations were modeled independently, the reconstruction propagated uncertainty from each fitted model without explicitly estimating cross-element posterior covariance. We also evaluated the physical plausibility of predicted elemental concentrations by identifying posterior combinations in which the sum of %C, %N, and %P exceeded 100%. These combinations were flagged for inspection rather than automatically excluded, preserving the original posterior distributions. The resulting predictions were used in the dietary reconstruction and subsequent sensitivity analyses.

**Figure S1. Estimated dietary composition of guppies across urbanization origins.** Contributions of four dietary resources to guppy diets in LU and HU sites. Large symbols show overall posterior medians and 95% credible intervals. Smaller symbols represent individual rivers, with lines connecting paired LU and HU sites. Annotations indicate the estimated HU−LU difference in percentage points and the posterior probability of the indicated direction. Estimates are based on the hierarchical Bayesian model with jointly estimated TDFs.

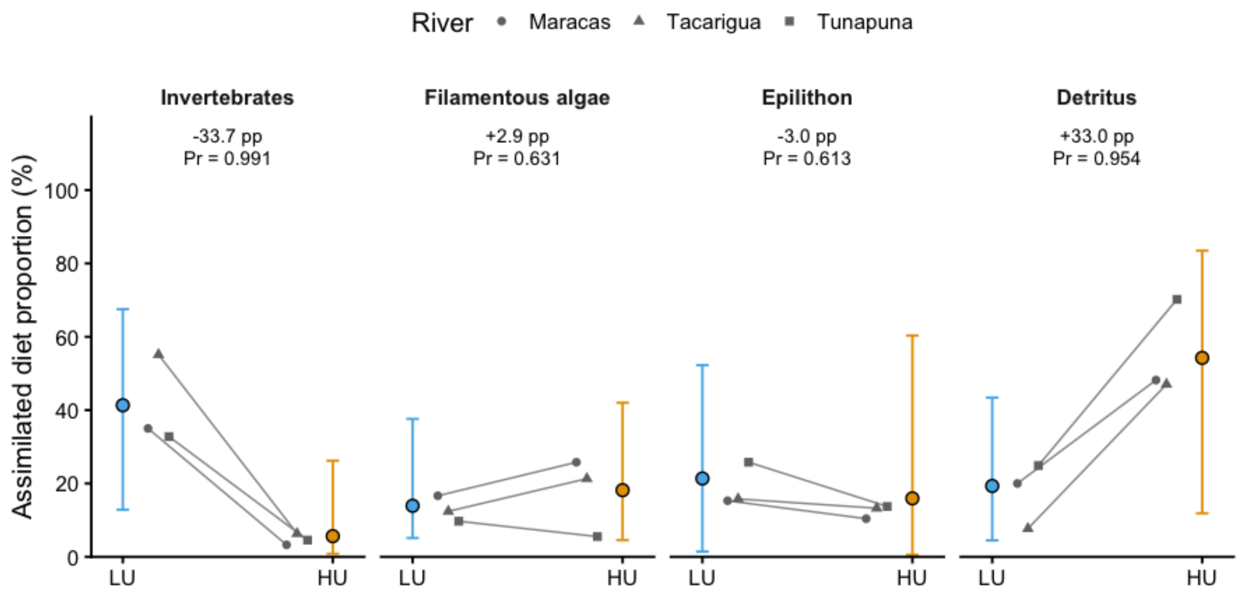

**Figure S2. Isotopic composition of guppies and dietary resources across urbanization origins.** $\delta^{13}C$ and $\delta^{15}N$ isotope values of guppies and four potential dietary resources across paired low-urbanization (LU) and high-urbanization (HU) sites in three rivers. Crosses represent individual guppies, small open circles represent measured resources, and large filled circles represent posterior medians of site-specific source isotope signatures, with horizontal and vertical bars indicating marginal 95% credible intervals. Guppy carbon values were lipid-normalized, guppy and resource nitrogen values were centered relative to site-specific snail baselines, and posterior median trophic discrimination factors (TDFs) were subtracted from guppy isotope values for visualization. The number of plotted resource observations reflects sampling effort and taxonomic coverage rather than resource abundance or dietary contribution.

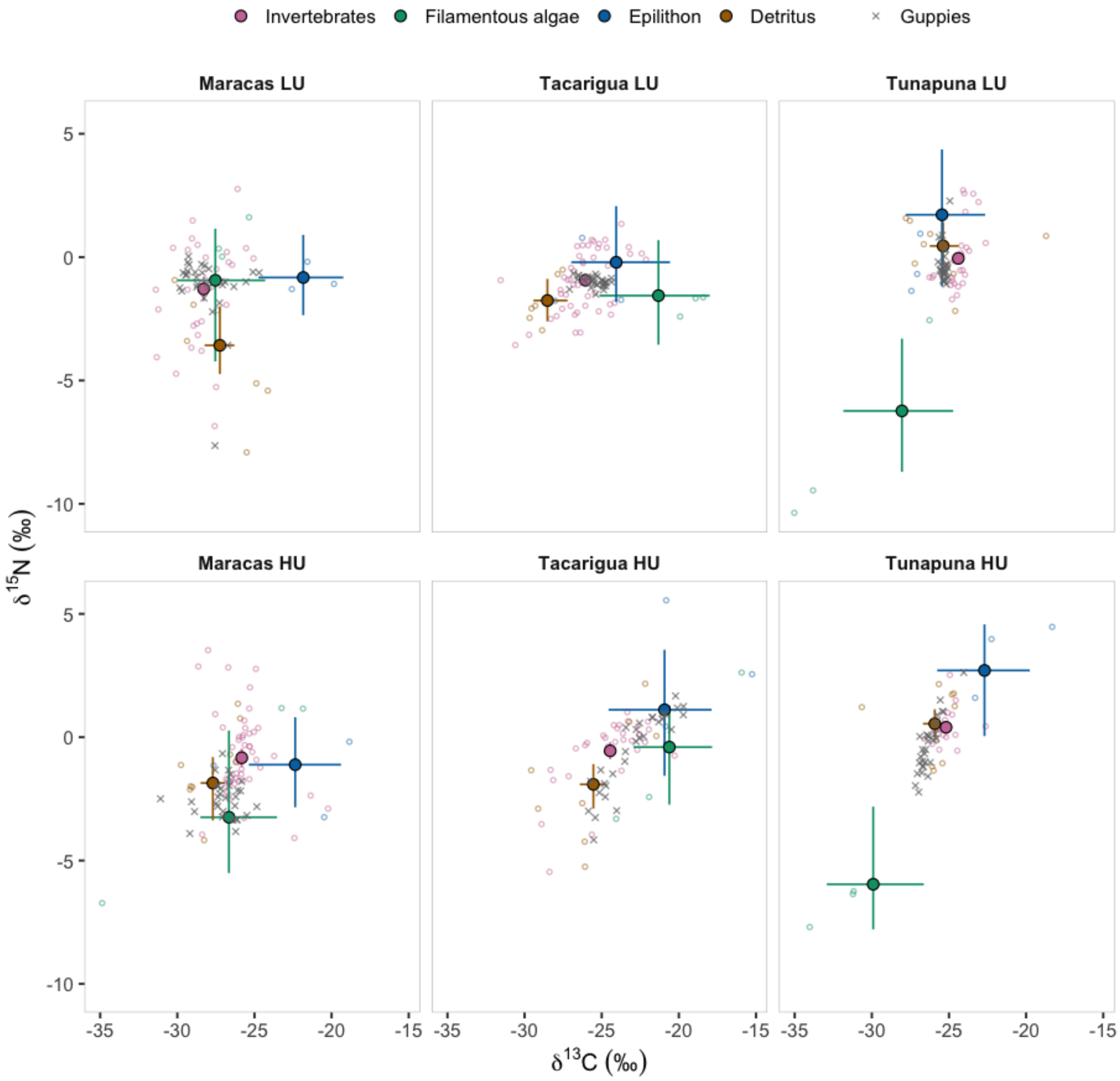

**Figure S3. Posterior correlations among dietary source contributions in the hierarchical Bayesian mixing model.** Pearson correlation coefficients (r) among the estimated dietary contributions of detritus, epilithon, filamentous algae, and aquatic invertebrates in low-urbanization (LU) and high-urbanization (HU) populations and their paired HU−LU contrasts. Correlations were calculated across 20,000 aligned posterior draws from the primary model with jointly estimated trophic discrimination factors. Origin-level contributions represent equal-weighted averages across the three rivers, and HU−LU contrasts represent equal-weighted averages of within-river differences, evaluated at reference somatic mass. Values within cells indicate Pearson correlation coefficients, with shading representing their direction and magnitude. Correlations describe dependence among posterior estimates of dietary proportions, which are constrained to sum to one, and do not independently establish ecological resource substitution or source identifiability.

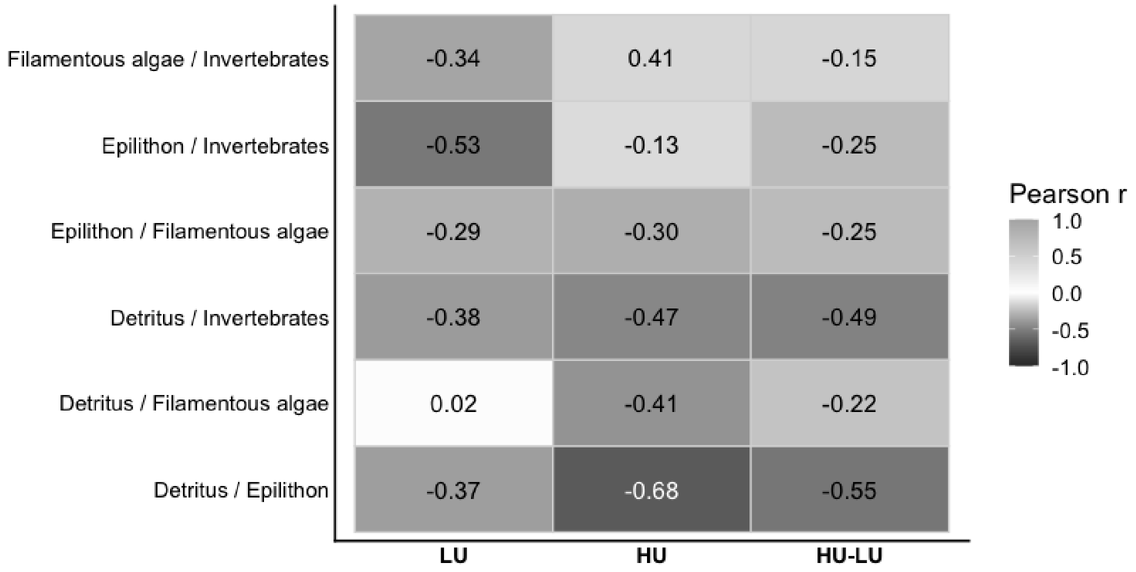

**Figure S4. Resource-specific associations between environmental gradients and stoichiometry.** Points represent observed log-transformed molar ratios, with circles and triangles indicating low-urbanization (LU) and high-urbanization (HU) sites, respectively. Lines show fitted mixed-effects model relationships, with shaded bands indicating uncertainty. Line types distinguish detrital subtypes (CPOM, FPOM, and sewage), while point color represents total chlorophyll concentration in the epilithon C:N panel.

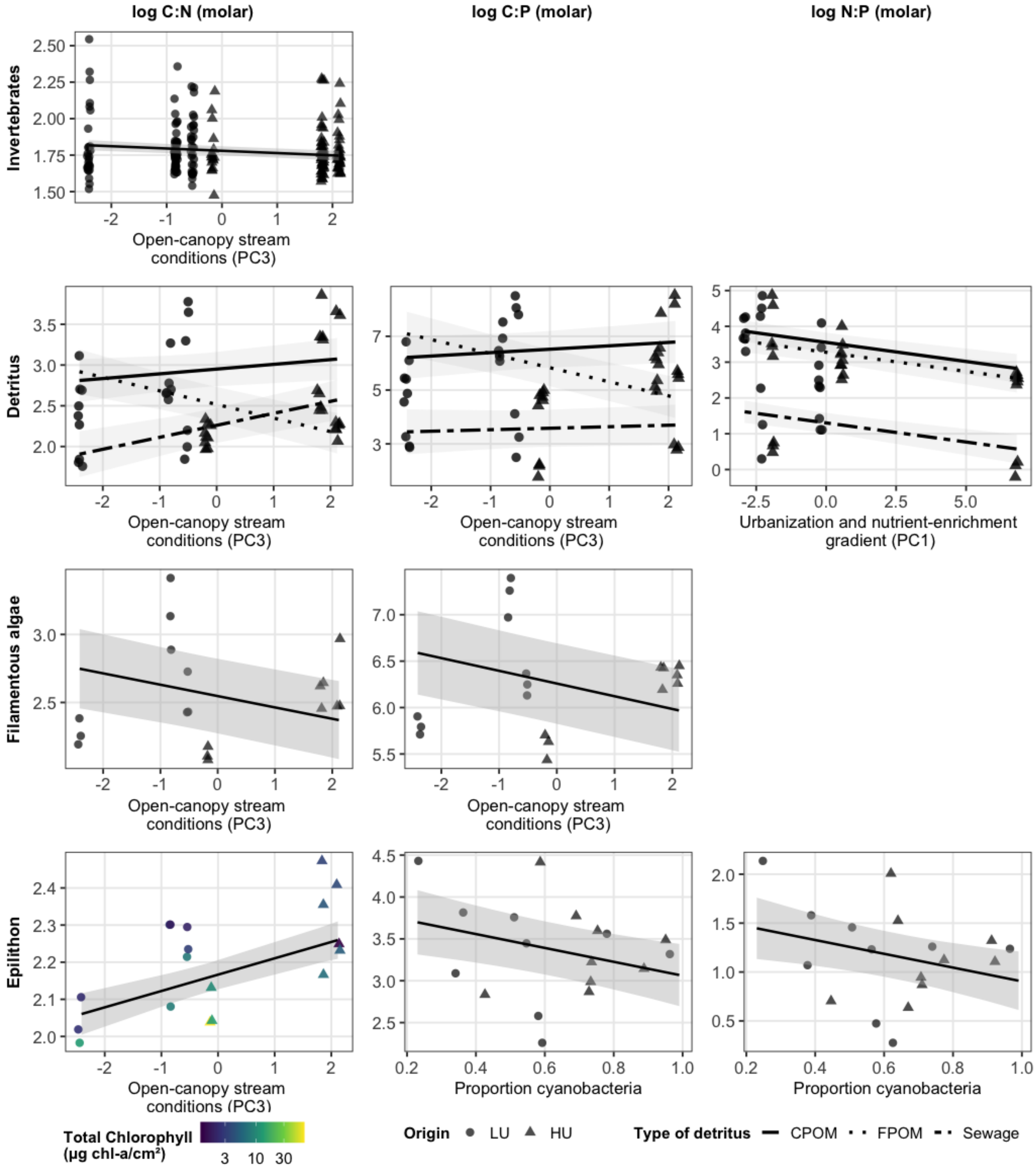

**Figure S5. Fish species presence-absence across paired urbanization sites.** Presence-absence matrix of fish species recorded at LU and HU sites in Maracas, Tacarigua, and Tunapuna rivers. Black squares indicate species presence and gray squares indicate absence. The matrix was used to calculate Jaccard dissimilarity for supplementary community-composition analyses.

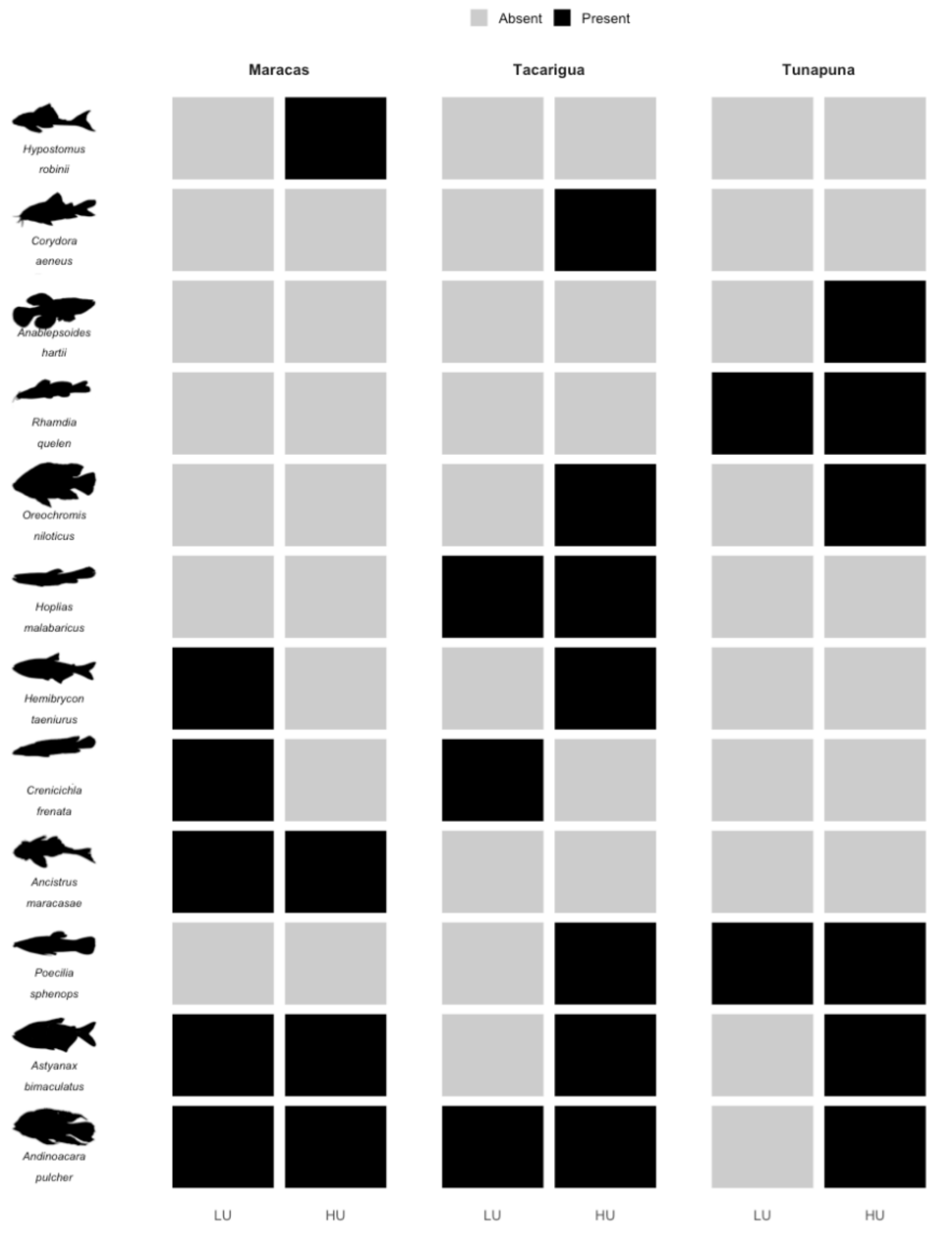

**Table S1. Fixed effects associated with basal-resource stoichiometry.**

| Resource | Ratio | Model formula | Coefficient reported | β ± SE | Test | p |
|---|---|---|---|---|---|---|
| **Epilithon** | C:N | log C:N ~ log(total chlorophyll) + PC1 + (1 \| River) + (1 \| Site) | PC1 | 0.020 ± 0.011 | t = 1.89, df = 13.95 | 0.080 |
| | | log C:N ~ log(total chlorophyll) + PC3 + (1 \| River) + (1 \| Site) | PC3 | 0.044 ± 0.015 | t = 3.01, df = 13.29 | **0.010** |
| | | | log(total chlorophyll) | -0.061 ± 0.024 | t = -2.57, df = 14.65 | **0.022** |
| | C:P | log C:P ~ cyanobacteria proportion + (1 \| River) + (1 \| Site) | Cyanobacteria proportion | -0.828 ± 0.493 | t = -1.68, df = 14.21 | 0.115 |
| | N:P | log N:P ~ cyanobacteria proportion + (1 \| River) + (1 \| Site) | Cyanobacteria proportion | -0.705 ± 0.476 | t = -1.48, df = 14.35 | 0.160 |
| **Filamentous algae** | C:N | log C:N ~ PC2 + (1 \| River) + (1 \| Site) | PC2 | 0.080 ± 0.055 | t = 1.45, df = 2.86 | 0.248 |
| | | log C:N ~ PC3 + (1 \| River) + (1 \| Site) | PC3 | -0.075 ± 0.056 | t = -1.32, df = 2.12 | 0.310 |
| | C:P | log C:P ~ PC2 + (1 \| River) + (1 \| Site) | PC2 | 0.135 ± 0.087 | t = 1.55, df = 2.91 | 0.222 |
| | | log C:P ~ PC3 + (1 \| River) + (1 \| Site) | PC3 | -0.111 ± 0.099 | t = -1.12, df = 2.15 | 0.372 |
| | N:P | log N:P ~ PC2 + (1 \| River) + (1 \| Site) | PC2 | 0.055 ± 0.032 | t = 1.7, df = 2.95 | 0.189 |
| | | log N:P ~ PC3 + (1 \| River) + (1 \| Site) | PC3 | -0.035 ± 0.043 | t = -0.83, df = 2.21 | 0.488 |
| **Invertebrates** | C:N | log C:N ~ PC3 + (1 \| River) + (1 \| Site) + (1 \| Taxa) | PC3 | -0.016 ± 0.007 | t = -2.29, df = 197.9 | **0.023** |
| | C:P | log C:P ~ PC3 + (1 \| River) + (1 \| Site) + (1 \| Taxa) | PC3 | -0.053 ± 0.035 | t = -1.5, df = 185.14 | 0.136 |
| | N:P | log N:P ~ PC3 + (1 \| River) + (1 \| Site) + (1 \| Taxa) | PC3 | -0.040 ± 0.035 | t = -1.14, df = 185.68 | 0.257 |
| **Detritus** | C:N | log C:N ~ PC1 × detritus type + (1 \| River) + (1 \| Site) | PC1 slope in CPOM | -0.097 ± 0.042 | t = -2.3, df = 5.92 | 0.062 |
| | | | PC1 slope in FPOM | -0.000 ± 0.045 | t = -0.01, df = 6.92 | 0.994 |
| | | | PC1 slope in Sewage | 0.026 ± 0.044 | t = 0.59, df = 6.11 | 0.579 |
| | | log C:N ~ PC3 × detritus type + (1 \| River) + (1 \| Site) | PC3 slope in CPOM | 0.060 ± 0.072 | t = 0.83, df = 9.43 | 0.426 |
| | | | PC3 slope in FPOM | -0.165 ± 0.071 | t = -2.32, df = 8.81 | **0.046** |
| | | | PC3 slope in Sewage | 0.149 ± 0.074 | t = 2.02, df = 9.71 | 0.072 |
| | C:P | log C:P ~ PC1 + detritus type + (1 \| River) + (1 \| Site) | FPOM vs CPOM | -0.717 ± 0.387 | t = -1.85, df = 41.45 | 0.071 |
| | | | PC1 | -0.129 ± 0.063 | t = -2.06, df = 32.6 | **0.048** |
| | | | Sewage vs CPOM | -2.901 ± 0.385 | t = -7.54, df = 41.17 | **<0.001** |
| | | log C:P ~ PC3 × detritus type + (1 \| River) + (1 \| Site) | PC3 slope in CPOM | 0.131 ± 0.183 | t = 0.72, df = 8.87 | 0.492 |
| | | | PC3 slope in FPOM | -0.519 ± 0.180 | t = -2.88, df = 8.26 | **0.020** |
| | | | PC3 slope in Sewage | 0.060 ± 0.187 | t = 0.32, df = 9.07 | 0.755 |
| | N:P | log N:P ~ PC1 + detritus type + (1 \| River) + (1 \| Site) | FPOM vs CPOM | -0.277 ± 0.266 | t = -1.04, df = 41.63 | 0.304 |
| | | | PC1 | -0.107 ± 0.042 | t = -2.56, df = 24.39 | **0.017** |
| | | | Sewage vs CPOM | -2.250 ± 0.265 | t = -8.49, df = 41.24 | **<0.001** |
| | | log N:P ~ PC3 + detritus type + (1 \| River) + (1 \| Site) | PC3 | -0.120 ± 0.087 | t = -1.37, df = 42.99 | 0.176 |

*Each row reports one fixed-effect coefficient or simple slope from the model formula shown. Rows sharing the same model formula come from the same fitted model. All stoichiometric ratios were log-transformed before analysis. Regression coefficients are reported as β ± SE on the log-response scale. PC1 represents nutrient enrichment, PC2 represents the nitrogen/benthic-habitat gradient, and PC3 represents open-canopy stream conditions. CPOM was the reference level for detritus contrasts. Detritus slopes from interaction models were estimated using emmeans::emtrends(). River identity was included as a random intercept in all models, with Site (River × Origin) as an additional random intercept; invertebrate models also included taxon identity as a random intercept. Bold p-values indicate p < 0.05; all PC associations are exploratory because only six sites were sampled.*

**Table S2. Prior distributions for the primary hierarchical isotope mixing model**

| Model component | JAGS parameter | Prior | Scale |
|---|---|---|---|
| Baseline dietary proportions | p.global | Dirichlet(2.033, 1.068, 1.652, 1.247) | Proportions |
| Urbanization effect | ilr.treat[2,k] | Normal(0, 1) | ILR |
| Somatic-mass effect | ilr.mass[k] | Normal(0, 0.5) | ILR per 1 SD log mass |
| Dietary river SD | sd.river | Half-normal(0, 0.50) | ILR SD |
| Dietary site SD | sd.diet.site | Half-normal(0, 0.35) | ILR SD |
| Overall source isotope means | src.global[s,j] | Normal(0, 5) | Standardized isotope |
| Source river SD | sd.src.river[j] | Half-normal(0, 0.70) | Standardized isotope SD |
| Source site SD | sd.src.site[j] | Half-normal(0, 0.70) | Standardized isotope SD |
| Detritus/invertebrate subgroup SD | sd.subgroup[1 or 4,j] | Half-normal(0, 0.70) | Standardized isotope SD |
| Epilithon/algal subgroup SD | sd.subgroup[2 or 3,j] | Fixed at zero | Not estimated |
| Within-subgroup analytical SD | sd.within.global[s,j] | Half-normal(0, 1.00) | Standardized isotope SD |
| Site variation in source log-SD | sd.within.spread[j] | Half-normal(0, 0.35) | Dimensionless log-SD |
| Consumer residual precision | Tau.base | Wishart(I, 3) | Standardized isotope precision |
| Site variation in residual log-SD | sd.log.res[j] | Half-normal(0, 0.50) | Dimensionless log-SD |
| Carbon trophic discrimination | tdf.latent[1] | Normal(0.40, 1.30) | ‰ |
| Nitrogen trophic discrimination | tdf.latent[2] | Normal(2.54, 1.27) | ‰ |
| Non-centered random-effect variables | *.raw terms | Independent Normal(0, 1) | Standard-normal latent |

*Normal and half-normal distributions are parameterized by mean and SD, not precision; half-normal distributions are restricted to nonnegative values. I denotes the 2 x 2 identity matrix used in the Wishart prior. Isotope-source scales refer to values after standardization. The carbon and nitrogen TDF priors are reported in per mil. Subgroup SDs for epilithon and filamentous algae are fixed at zero because these sources have no modeled subgroups.*

**Table S3. Primary model: assimilated dietary contributions by origin**

| Dietary source | LU contribution (%) | HU contribution (%) | HU − LU (pp) | *Pr*(HU > LU) |
|---|---|---|---|---|
| Detritus | 19.3 [4.5, 43.4] | 54.2 [11.9, 83.5] | +33.0 [-4.6, +64.2] | 0.954 |
| Epilithon | 21.4 [1.4, 52.3] | 15.9 [0.5, 60.4] | -3.0 [-32.3, +33.4] | 0.387 |
| Filamentous algae | 13.9 [5.1, 37.6] | 18.2 [4.6, 42.0] | +2.9 [-15.4, +25.0] | 0.631 |
| Invertebrates | 41.3 [12.8, 67.5] | 5.6 [0.8, 26.2] | -33.7 [-60.1, -5.0] | 0.009 |

*Primary specification jointly estimates globally shared carbon and nitrogen TDFs. Origin values summarize site-level contributions averaged equally across the three rivers. The HU−LU contrast is computed within each river and then averaged; it is not the difference of independently reported posterior medians. pp = percentage points. Because each source is summarized using its marginal posterior median, the reported median contributions do not necessarily sum to 100%, although the proportions sum to one within each posterior draw.*

**Table S4. Primary model: paired HU−LU contrasts within each river**

| Dietary source | Maracas (pp) | Tacarigua (pp) | Tunapuna (pp) |
|---|---|---|---|
| Detritus | +25.7 [-21.1, +72.2] | +36.0 [-3.2, +68.7] | +37.3 [-11.3, +76.3] |
| Epilithon | -2.4 [-37.0, +38.5] | -0.4 [-41.2, +56.3] | -5.3 [-48.1, +37.3] |
| Filamentous algae | +4.4 [-29.2, +55.9] | +7.0 [-23.4, +35.4] | -3.8 [-14.6, +6.0] |
| Invertebrates | -30.4 [-72.2, -1.3] | -44.1 [-80.5, -1.7] | -24.8 [-55.8, +5.3] |

*Posterior medians [95% CrI] of river-specific HU−LU site contrasts at reference somatic mass; estimates are partially pooled through the model hierarchy.*

**Table S5. TDF sensitivity: pooled within-river dietary contrasts**

| TDF specification | Dietary source | HU − LU (pp) | *Pr*(HU > LU) |
|---|---|---|---|
| Estimated TDF (primary) | Detritus | +33.0 [-4.6, +64.2] | 0.954 |
| Estimated TDF (primary) | Epilithon | -3.0 [-32.3, +33.4] | 0.387 |
| Estimated TDF (primary) | Filamentous algae | +2.9 [-15.4, +25.0] | 0.631 |
| Estimated TDF (primary) | Invertebrates | -33.7 [-60.1, -5.0] | 0.009 |
| Conventional fixed | Detritus | -2.0 [-22.5, +17.6] | 0.368 |
| Conventional fixed | Epilithon | +12.1 [-21.8, +46.0] | 0.774 |
| Conventional fixed | Filamentous algae | -6.3 [-34.8, +24.9] | 0.311 |
| Conventional fixed | Invertebrates | -2.6 [-19.3, +5.6] | 0.211 |
| Stephens C3 | Detritus | -2.3 [-29.1, +22.2] | 0.399 |
| Stephens C3 | Epilithon | +2.5 [-26.0, +31.0] | 0.586 |
| Stephens C3 | Filamentous algae | +3.8 [-18.7, +31.4] | 0.629 |
| Stephens C3 | Invertebrates | -4.1 [-15.1, +4.5] | 0.128 |
| Stephens mixed | Detritus | +6.7 [-17.5, +35.1] | 0.698 |
| Stephens mixed | Epilithon | +2.9 [-30.3, +36.4] | 0.577 |
| Stephens mixed | Filamentous algae | +3.4 [-15.4, +28.6] | 0.631 |
| Stephens mixed | Invertebrates | -14.5 [-33.0, +2.2] | 0.039 |

*All scenarios use identical non-TDF data and model settings. The estimated-TDF specification is the selected primary analysis. Directional probabilities refer to posterior support for positive HU−LU contrasts; for negative contrasts, use 1 − Pr(HU > LU).*

**Table S6. Sensitivity of reconstructed dietary stoichiometry and its decomposition to trophic discrimination factors.**

| TDF specification | Ratio | HU − LU (%) | Pr(HU > LU) | Pr(\|C\| > \|Q\|) |
|---|---|---|---|---|
| Estimated (C 2.82, N 3.00; primary) | C:N | +42.7 [-9.0, +132.6] | 0.940 | 0.935 |
| Estimated (C 2.82, N 3.00; primary) | C:P | -41.3 [-69.7, +10.5] | 0.046 | 0.472 |
| Estimated (C 2.82, N 3.00; primary) | N:P | -59.2 [-81.8, -10.0] | 0.013 | 0.908 |
| Conventional fixed (C 0.40, N 2.54) | C:N | -7.5 [-39.2, +40.4] | 0.349 | 0.355 |
| Conventional fixed (C 0.40, N 2.54) | C:P | -38.7 [-74.3, +26.1] | 0.091 | 0.552 |
| Conventional fixed (C 0.40, N 2.54) | N:P | -34.2 [-73.5, +47.3] | 0.157 | 0.694 |
| Stephens C3 (C 1.20, N 4.10) | C:N | -5.5 [-36.0, +37.6] | 0.382 | 0.345 |
| Stephens C3 (C 1.20, N 4.10) | C:P | -28.0 [-59.4, +30.6] | 0.126 | 0.312 |
| Stephens C3 (C 1.20, N 4.10) | N:P | -24.1 [-60.3, +49.3] | 0.201 | 0.527 |
| Stephens mixed (C 1.40, N 3.10) | C:N | +15.7 [-20.8, +70.7] | 0.781 | 0.768 |
| Stephens mixed (C 1.40, N 3.10) | C:P | -34.0 [-66.9, +25.6] | 0.092 | 0.466 |
| Stephens mixed (C 1.40, N 3.10) | N:P | -43.6 [-74.4, +20.7] | 0.069 | 0.785 |

*Values are posterior medians [95% credible intervals] of HU − LU percentage changes in reconstructed dietary molar ratios, calculated within each river and averaged across the three paired rivers. Pr(HU > LU) is the posterior probability that the contrast is positive. Pr(|C| > |Q|) is the probability that the absolute diet-composition contribution exceeds the resource-quality contribution on the log-ratio scale. Each row reconstructs dietary stoichiometry from the dietary posterior of the indicated mixing-model specification. TDF values are carbon and nitrogen discrimination factors (‰); for the estimated specification they are posterior medians.*

**Table S7. Literature-informed TDF priors and estimated posterior**

| Isotope | Normal prior: mean ± SD (‰) | Posterior median [95% CrI] (‰) |
|---|---|---|
| Carbon ($\delta^{13}C$) | 0.40 ± 1.30 | 2.82 [2.20, 3.33] |
| Nitrogen ($\delta^{15}N$) | 2.54 ± 1.27 | 3.00 [2.72, 3.41] |

*The literature-derived prior SD characterizes the prior for a globally shared parameter; it is not an estimate of within-population guppy TDF variation. Posterior intervals are conditional on the mixing model and its source assumptions.*

**Table S8. Posterior TDF–diet dependence and sampling diagnostics under the primary model**

| Posterior parameter | r with C TDF | r with N TDF | $\hat{R}$ | Bulk ESS | Tail ESS |
|---|---|---|---|---|---|
| Detritus contrast | +0.190 | −0.164 | 1.0002 | 11,111 | 15,387 |
| Epilithon contrast | −0.027 | +0.028 | 1.0003 | 11,027 | 14,997 |
| Filamentous-algae contrast | −0.067 | −0.070 | 1.0006 | 9,777 | 11,125 |
| Invertebrate contrast | −0.163 | +0.224 | 1.0008 | 5,660 | 9,599 |
| Carbon TDF | — | — | 1.0003 | 7,856 | 7,698 |
| Nitrogen TDF | — | — | 1.0003 | 9,170 | 7,490 |

*Pearson correlations quantify within-model associations between each TDF and the paired HU−LU dietary contrast for the indicated source. The contrasts are equal-river averages in percentage points. $\hat{R}$, bulk ESS, and tail ESS summarize posterior sampling across the completed chains. Weak linear correlations do not establish TDF–diet independence or remove sensitivity to alternative fixed-TDF specifications.*

**Table S9. Source-only versus primary joint-model source signatures**

| Source | Isotope | Median \|shift\| (‰) | Maximum \|shift\| (‰) | Outside SO 95% CrI |
|---|---|---|---|---|
| Detritus | $\delta^{13}C$ | 0.06 | 0.13 | 0/6 |
| Detritus | $\delta^{15}N$ | 0.22 | 0.36 | 0/6 |
| Epilithon | $\delta^{13}C$ | 0.08 | 0.37 | 0/6 |
| Epilithon | $\delta^{15}N$ | 0.08 | 1.36 | 0/6 |
| Filamentous algae | $\delta^{13}C$ | 0.68 | 1.75 | 0/6 |
| Filamentous algae | $\delta^{15}N$ | 0.46 | 2.12 | 0/6 |
| Invertebrates | $\delta^{13}C$ | 0.01 | 0.04 | 0/6 |
| Invertebrates | $\delta^{15}N$ | 0.03 | 0.11 | 0/6 |

*Shift = posterior median from the joint primary model minus the source-only posterior median for the same site, source, and isotope. Six site comparisons per row; there were 0/48 joint-model medians outside source-only 95% CrIs. Interval overlap is descriptive and not a formal equivalence test.*

**Table S10. Source-estimate displacement across trophic discrimination specifications**

| TDF specification | Mean absolute shift (‰) | Maximum shift (‰) | Outside source-only 95% CrI |
|---|---|---|---|
| Jointly estimated (primary) | 0.319 | 2.124 | 0/48 |
| Conventional fixed | 0.965 | 7.734 | 4/48 |
| Stephens C3 | 0.694 | 6.927 | 3/48 |
| Stephens mixed | 0.700 | 7.115 | 2/48 |

*Shifts are absolute differences between fitted source-isotope posterior medians and the corresponding source-only estimates across 48 site–source–isotope comparisons. The counts describe joint-model medians outside the source-only 95% credible intervals. These comparisons are diagnostic, not model-selection scores.*

**Table S11. Final-run computational and posterior predictive diagnostics**

| Model | Maximum core $\hat{R}$ | Minimum core bulk ESS | Consumer PPC flags | Source PPC flags |
|---|---|---|---|---|
| Estimated TDF (primary) | 1.004 | 1,026 | 0/12 | 0/48 |
| Conventional fixed TDF | 1.002 | 1,695 | 0/12 | 0/48 |
| Stephens C3 | 1.005 | 2,297 | 0/12 | 0/48 |
| Stephens mixed | 1.002 | 1,594 | 0/12 | 0/48 |
| Symmetric urbanization coding | 1.005 | 847 | 0/12 | 0/48 |
| Wider dietary site-SD prior (0.70) | 1.003 | 977 | 0/12 | 0/48 |
| Flat Dirichlet(1, 1, 1, 1) prior | 1.005 | 862 | 0/12 | 0/48 |

*All seven completed mixing-model specifications were fitted using four MCMC chains of 1,000,000 iterations each, including 500,000 burn-in iterations and thinning by 100. Maximum core $\hat{R}$ and minimum core bulk effective sample size (ESS) summarize monitored core parameters. Posterior predictive check (PPC) flags denote RMSE tail probabilities below 0.025 or above 0.975; consumer checks cover 12 site–isotope combinations and source checks cover 48 site–source–isotope combinations. The source-only diagnostic model (not shown) had maximum $\hat{R}$ = 1.008 and minimum bulk ESS = 920. Satisfactory diagnostics do not establish unique dietary-mixture identifiability.*

**Table S12. Sensitivity of paired HU−LU dietary contrasts to urbanization-prior parameterization**

| Dietary source | LU-reference HU−LU (pp) | Symmetric-origin HU−LU (pp) |
|---|---|---|
| Detritus | +33.0 [−4.6, 64.2] | +37.2 [1.3, 67.3] |
| Epilithon | −3.0 [−32.3, 33.4] | −3.7 [−37.2, 29.5] |
| Filamentous algae | +2.9 [−15.4, 25.0] | +5.5 [−12.1, 26.4] |
| Invertebrates | −33.7 [−60.1, −5.0] | −38.9 [−65.0, −7.2] |

*Posterior medians [95% CrI] are equal-weighted averages of paired HU−LU contrasts across three rivers, evaluated at reference somatic mass. Both specifications use jointly estimated TDFs, identical data and non-origin prior settings, and a Normal(0, 1) prior on the HU−LU ILR coefficient. The symmetric specification changes the marginal priors induced for LU and HU dietary proportions. Convergence diagnostics for the symmetric specification were satisfactory (maximum core $\hat{R}$ = 1.005; minimum core bulk ESS = 847; minimum core tail ESS = 1,718).*

**Table S13. Sensitivity of paired HU−LU dietary contrasts to the dietary site-effect prior scale**

| Dietary source | Primary HU−LU (pp) | ***Pr*(HU > LU)** | Wider-prior HU−LU (pp) | ***Pr*(HU > LU)** |
|---|---|---|---|---|
| Detritus | +33.0 [−4.6, 64.2] | 0.954 | +33.6 [−4.2, 68.0] | 0.958 |
| Epilithon | −3.0 [−32.3, 33.4] | 0.387 | −2.1 [−34.3, 36.9] | 0.420 |
| Filamentous algae | +2.9 [−15.4, 25.0] | 0.631 | +6.9 [−15.7, 32.2] | 0.730 |
| Invertebrates | −33.7 [−60.1, −5.0] | 0.009 | −40.1 [−67.4, −7.1] | 0.008 |

*Values are posterior medians [95% credible intervals] of HU−LU differences in assimilated dietary contributions, expressed in percentage points (pp). Contrasts were calculated within each river at reference somatic mass and averaged equally across the three paired rivers. Pr(HU > LU) represents the posterior probability of a positive contrast. The two models differ only in the half-normal prior scale assigned to the dietary site-effect standard deviation (0.35 versus 0.70 on the ILR scale); both retain jointly estimated trophic discrimination factors and identical remaining model specifications.*

**Table S14. Flat versus gut-informed global dietary prior**

| **Dietary source** | **Gut prior HU-LU (pp)** | ***Pr*(HU>LU), gut** | **Flat prior HU-LU (pp)** | ***Pr*(HU>LU), flat** | **Median change (pp)** |
|---|---|---|---|---|---|
| Detritus | +33.0 [−4.6, 64.2] | 0.954 | +32.6 [-3.4, +64.7] | 0.953 | -0.41 |
| Epilithon | −3.0 [−32.3, 33.4] | 0.387 | -3.9 [-35.3, +32.0] | 0.382 | -0.81 |
| Filamentous algae | +2.9 [−15.4, 25.0] | 0.631 | +4.1 [-12.2, +26.5] | 0.696 | +1.16 |
| Invertebrates | −33.7 [−60.1, −5.0] | 0.009 | -34.0 [-60.7, -4.0] | 0.011 | -0.27 |

*Values are posterior medians [95% credible intervals] of HU−LU differences in assimilated dietary contributions (pp), calculated within each river at reference somatic mass and averaged equally across the three rivers. The flat Dirichlet(1, 1, 1, 1) prior replaces only the gut-informed baseline dietary proportions; all other hierarchical dietary priors are unchanged.*

**Table S15. Sensitivity of reconstructed dietary stoichiometry and its decomposition to resource elemental prediction scale.**

| Scenario | Ratio | LU | HU | Total | Diet composition | Resource quality | Pr(HU > LU) | Pr(\|C\| > \|Q\|) |
|---|---|---|---|---|---|---|---|---|
| Primary | C:N | 7.4<br>[5.9, 11.1] | 10.8<br>[7.2, 17.3] | +42.7<br>[-9.0, +132.6] | +54.5<br>[+7.4, +134.9] | -7.7<br>[-29.2, +17.8] | 0.940 | 0.935 |
| Primary | C:P | 445.8<br>[254.3, 685.2] | 260.2<br>[127.6, 468.1] | -41.3<br>[-69.7, +10.5] | -21.4<br>[-57.6, +31.5] | -24.9<br>[-46.4, +5.3] | 0.046 | 0.472 |
| Primary | N:P | 59.2<br>[29.6, 97.4] | 23.7<br>[10.9, 48.2] | -59.2<br>[-81.8, -10.0] | -49.8<br>[-76.0, -1.5] | -18.3<br>[-42.7, +17.3] | 0.013 | 0.908 |
| Conditional median | C:N | 7.6<br>[6.0, 11.5] | 11.7<br>[7.6, 19.1] | +50.7<br>[-6.4, +150.8] | +63.1<br>[+8.9, +153.8] | -7.4<br>[-28.4, +17.7] | 0.956 | 0.954 |
| Conditional median | C:P | 502.5<br>[280.3, 777.7] | 289.0<br>[140.2, 519.4] | -42.2<br>[-70.4, +10.1] | -22.6<br>[-59.0, +30.9] | -24.7<br>[-46.2, +5.8] | 0.046 | 0.499 |
| Conditional median | N:P | 65.2<br>[31.3, 109.4] | 24.2<br>[11.0, 50.7] | -62.2<br>[-83.5, -12.6] | -53.3<br>[-78.2, -3.6] | -18.5<br>[-42.4, +16.6] | 0.012 | 0.924 |

*Entries for LU and HU are posterior medians [95% credible intervals] of absolute dietary molar ratios (mol:mol). Total, diet composition, and resource quality are HU–LU percentage changes [95% credible intervals]. Pr(HU > LU) is the posterior probability that the total contrast is positive. Pr(|C| > |Q|) is the probability that the absolute diet-composition contribution exceeds the resource-quality contribution on the log-ratio scale. Components sum to the total on the log scale, not after conversion to percentages. All scenarios use the same 10,000 paired draws from the primary dietary reconstruction, which is repeated in each table for reference. The primary reconstruction used conditional arithmetic mean resource elemental concentrations; the alternative used conditional medians. Detrital %C, fitted with beta regression, used the inverse-logit fitted mean in both cases.*

**Table S16. Sensitivity of reconstructed dietary stoichiometry and its decomposition to detrital-subtype weighting.**

<table>
<tr><th>Scenario</th><th>Ratio</th><th>LU</th><th>HU</th><th>Total</th><th>Diet composition</th><th>Resource quality</th><th>Pr(HU > LU)</th><th>Pr(|C| > |Q|)</th></tr>
<tr><td>Primary</td><td>C:N</td><td>7.4 [5.9, 11.1]</td><td>10.8 [7.2, 17.3]</td><td>+42.7 [-9.0, +132.6]</td><td>+54.5 [+7.4, +134.9]</td><td>-7.7 [-29.2, +17.8]</td><td>0.940</td><td>0.935</td></tr>
<tr><td>Primary</td><td>C:P</td><td>445.8 [254.3, 685.2]</td><td>260.2 [127.6, 468.1]</td><td>-41.3 [-69.7, +10.5]</td><td>-21.4 [-57.6, +31.5]</td><td>-24.9 [-46.4, +5.3]</td><td>0.046</td><td>0.472</td></tr>
<tr><td>Primary</td><td>N:P</td><td>59.2 [29.6, 97.4]</td><td>23.7 [10.9, 48.2]</td><td>-59.2 [-81.8, -10.0]</td><td>-49.8 [-76.0, -1.5]</td><td>-18.3 [-42.7, +17.3]</td><td>0.013</td><td>0.908</td></tr>
<tr><td>Site occurrence</td><td>C:N</td><td>7.5 [5.9, 11.2]</td><td>10.8 [7.2, 17.3]</td><td>+41.5 [-9.8, +130.5]</td><td>+55.5 [+7.5, +137.0]</td><td>-9.0 [-30.2, +16.1]</td><td>0.936</td><td>0.933</td></tr>
<tr><td>Site occurrence</td><td>C:P</td><td>453.9 [259.9, 696.1]</td><td>260.2 [127.6, 468.1]</td><td>-42.4 [-70.3, +8.2]</td><td>-19.6 [-56.6, +33.9]</td><td>-27.8 [-48.8, +1.2]</td><td>0.042</td><td>0.399</td></tr>
<tr><td>Site occurrence</td><td>N:P</td><td>59.7 [30.2, 97.9]</td><td>23.7 [10.9, 48.2]</td><td>-59.6 [-81.9, -10.9]</td><td>-49.0 [-75.5, -0.6]</td><td>-20.4 [-44.5, +14.5]</td><td>0.012</td><td>0.891</td></tr>
<tr><td>No sewage</td><td>C:N</td><td>7.6 [6.0, 11.3]</td><td>10.8 [7.1, 17.5]</td><td>+40.9 [-10.6, +132.8]</td><td>+53.2 [+6.8, +132.6]</td><td>-8.1 [-30.6, +19.1]</td><td>0.930</td><td>0.924</td></tr>
<tr><td>No sewage</td><td>C:P</td><td>476.9 [271.4, 727.9]</td><td>328.0 [159.6, 586.7]</td><td>-31.0 [-64.3, +30.4]</td><td>-9.0 [-51.5, +53.7]</td><td>-23.7 [-45.4, +6.9]</td><td>0.115</td><td>0.378</td></tr>
<tr><td>No sewage</td><td>N:P</td><td>62.1 [32.2, 101.1]</td><td>29.9 [13.9, 56.7]</td><td>-51.3 [-77.6, +1.7]</td><td>-41.1 [-71.2, +7.0]</td><td>-16.6 [-41.7, +19.8]</td><td>0.028</td><td>0.843</td></tr>
</table>

*Entries for LU and HU are posterior medians [95% credible intervals] of absolute dietary molar ratios (mol:mol). Total, diet composition, and resource quality are HU–LU percentage changes [95% credible intervals]. Pr(HU > LU) is the posterior probability that the total contrast is positive. Pr(|C| > |Q|) is the probability that the absolute diet-composition contribution exceeds the resource-quality contribution on the log-ratio scale. Components sum to the total on the log scale, not after conversion to percentages. All scenarios use the same 10,000 paired draws from the primary dietary reconstruction, which is repeated in each table for reference. The primary reconstruction assigned equal weights to CPOM, FPOM, and sewage-derived material. The detritus site-occurrence scenario used site-level analytical occurrence weights; the no-sewage scenario excluded sewage and weighted CPOM and FPOM equally. Occurrence weights are not measurements of environmental availability.*

**Table S17. Sensitivity of reconstructed dietary stoichiometry and its decomposition to hypothetical sewage contributions within detrital resources.**

| Scenario | Ratio | LU | HU | Total | Diet composition | Resource quality | Pr(HU > LU) | Pr($\lvert C \rvert > \lvert Q \rvert$) |
|---|---|---|---|---|---|---|---|---|
| Primary | C:N | 7.4 [5.9, 11.1] | 10.8 [7.2, 17.3] | +42.7 [-9.0, +132.6] | +54.5 [+7.4, +134.9] | -7.7 [-29.2, +17.8] | 0.940 | 0.935 |
| Primary | C:P | 445.8 [254.3, 685.2] | 260.2 [127.6, 468.1] | -41.3 [-69.7, +10.5] | -21.4 [-57.6, +31.5] | -24.9 [-46.4, +5.3] | 0.046 | 0.472 |
| Primary | N:P | 59.2 [29.6, 97.4] | 23.7 [10.9, 48.2] | -59.2 [-81.8, -10.0] | -49.8 [-76.0, -1.5] | -18.3 [-42.7, +17.3] | 0.013 | 0.908 |
| HU 50% / LU 10% | C:N | 7.6 [6.0, 11.3] | 10.8 [7.2, 17.3] | +39.7 [-10.4, +126.4] | +55.9 [+7.6, +137.5] | -10.2 [-30.9, +15.8] | 0.932 | 0.929 |
| HU 50% / LU 10% | C:P | 471.3 [269.6, 719.8] | 230.4 [112.1, 429.5] | -50.8 [-75.1, -5.2] | -19.6 [-56.6, +33.8] | -37.9 [-58.4, -10.7] | 0.016 | 0.232 |
| HU 50% / LU 10% | N:P | 61.3 [31.6, 99.7] | 20.9 [9.3, 45.1] | -65.2 [-84.8, -19.8] | -49.1 [-75.6, -0.7] | -30.8 [-54.1, +1.3] | 0.007 | 0.796 |
| HU 70% / LU 5% | C:N | 7.6 [6.0, 11.4] | 10.8 [7.2, 17.3] | +39.3 [-10.8, +126.6] | +56.7 [+7.8, +138.9] | -10.9 [-31.4, +15.9] | 0.931 | 0.927 |
| HU 70% / LU 5% | C:P | 476.2 [271.6, 726.5] | 197.4 [93.4, 384.3] | -58.2 [-79.8, -15.9] | -22.9 [-58.5, +29.4] | -44.8 [-65.3, -16.9] | 0.008 | 0.172 |
| HU 70% / LU 5% | N:P | 61.7 [31.9, 100.4] | 17.8 [7.6, 42.3] | -70.5 [-87.7, -25.9] | -51.4 [-77.0, -2.4] | -38.0 [-61.5, -5.8] | 0.005 | 0.740 |
| HU 70% / LU 0% | C:N | 7.6 [6.0, 11.4] | 10.8 [7.2, 17.3] | +39.0 [-11.0, +126.3] | +56.6 [+7.7, +138.9] | -11.1 [-31.7, +15.8] | 0.930 | 0.926 |
| HU 70% / LU 0% | C:P | 481.1 [274.1, 734.0] | 197.4 [93.4, 384.3] | -58.6 [-80.1, -16.6] | -22.0 [-58.0, +30.6] | -45.9 [-66.4, -17.6] | 0.007 | 0.152 |
| HU 70% / LU 0% | N:P | 62.2 [32.3, 101.1] | 17.8 [7.6, 42.3] | -70.7 [-87.8, -26.3] | -50.8 [-76.7, -2.0] | -39.1 [-62.5, -6.6] | 0.005 | 0.712 |

Entries for LU and HU are posterior medians [95% credible intervals] of absolute dietary molar ratios (mol:mol). Total, diet composition, and resource quality are HU–LU percentage changes [95% credible intervals]. Pr(HU > LU) is the posterior probability that the total contrast is positive. Pr(|C| > |Q|) is the probability that the absolute diet-composition contribution exceeds the resource-quality contribution on the log-ratio scale. Components sum to the total on the log scale, not after conversion to percentages. All scenarios use the same 10,000 paired draws from the primary dietary reconstruction, which is repeated in each table for reference. The primary reconstruction used equal weighting of CPOM, FPOM, and sewage-derived material. Hypothetical scenarios assigned sewage 50% in HU versus 10% in LU, 70% versus 5%, or 70% versus 0%, with remaining detritus contributions weighted between CPOM and FPOM according to site occurrence. These are assumptions, not measured contributions.

**Table S18. Sensitivity of reconstructed dietary stoichiometry and its decomposition to macroinvertebrate taxon weighting.**

| Scenario | Ratio | LU | HU | Total | Diet composition | Resource quality | Pr(HU > LU) | Pr(\|C\| > \|Q\|) |
|---|---|---|---|---|---|---|---|---|
| Primary | C:N | 7.4 [5.9, 11.1] | 10.8 [7.2, 17.3] | +42.7 [-9.0, +132.6] | +54.5 [+7.4, +134.9] | -7.7 [-29.2, +17.8] | 0.940 | 0.935 |
| Primary | C:P | 445.8 [254.3, 685.2] | 260.2 [127.6, 468.1] | -41.3 [-69.7, +10.5] | -21.4 [-57.6, +31.5] | -24.9 [-46.4, +5.3] | 0.046 | 0.472 |
| Primary | N:P | 59.2 [29.6, 97.4] | 23.7 [10.9, 48.2] | -59.2 [-81.8, -10.0] | -49.8 [-76.0, -1.5] | -18.3 [-42.7, +17.3] | 0.013 | 0.908 |
| Site occurrence | C:N | 7.4 [5.9, 11.0] | 10.8 [7.1, 17.3] | +43.2 [-8.7, +133.5] | +55.2 [+7.5, +136.1] | -7.8 [-29.1, +17.8] | 0.940 | 0.936 |
| Site occurrence | C:P | 402.9 [236.6, 614.2] | 250.8 [125.4, 439.7] | -37.7 [-67.2, +14.4] | -12.0 [-51.8, +43.9] | -28.8 [-48.4, -0.2] | 0.058 | 0.296 |
| Site occurrence | N:P | 53.7 [28.3, 86.8] | 22.9 [10.8, 44.1] | -56.8 [-80.2, -10.7] | -43.9 [-72.5, +2.7] | -22.5 [-45.1, +11.3] | 0.012 | 0.826 |

*Entries for LU and HU are posterior medians [95% credible intervals] of absolute dietary molar ratios (mol:mol). Total, diet composition, and resource quality are HU–LU percentage changes [95% credible intervals]. Pr(HU > LU) is the posterior probability that the total contrast is positive. Pr(|C| > |Q|) is the probability that the absolute diet-composition contribution exceeds the resource-quality contribution on the log-ratio scale. Components sum to the total on the log scale, not after conversion to percentages. All scenarios use the same 10,000 paired draws from the primary dietary reconstruction, which is repeated in each table for reference. The primary reconstruction assigned equal weights to the 26 modeled macroinvertebrate taxa. The alternative used site-occurrence weighting of taxa recorded in analytical samples, not measured taxon abundance or ecological absence.*

**Table S19. Frequency of occurrence of gut-content categories in guppies sampled at each site.**

| River | Origin | Guts examined | Detritus (%) | Filamentous Algae (%) | Invertebrates (%) |
|---|---|---|---|---|---|
| Maracas | LU | 5 | 100 | 20 | 0 |
| Maracas | HU | 12 | 91.7 | 33.3 | 8.3 |
| Tacarigua | LU | 5 | 20 | 100 | 0 |
| Tacarigua | HU | 11 | 81.8 | 72.7 | 100 |
| Tunapuna | LU | – | – | – | – |
| Tunapuna | HU | 5 | 100 | 0 | 0 |

*Gut content categories represent the frequency of occurrence among examined fish at each site, calculated as the percentage of fish in which a given category was recorded. Detritus includes fish in which either detritus or sewage was recorded; filamentous algae includes fish in which filamentous algae was recorded; and invertebrates includes fish in which any of the following was recorded: Chironomus, Leptohyphidae, or snails. For each fish, a category was scored as present (1) if at least one of its constituent items was recorded and absent (0) otherwise. Categories are not mutually exclusive because multiple categories could occur in the same fish; consequently, percentages do not necessarily sum to 100%. Dashes indicate that no gut-content samples were available for Tunapuna LU. For the gut-informed Dirichlet prior, site-level occurrence frequencies were first averaged equally across sampled sites within each origin, and HU and LU means were then averaged equally. Thus, sites were weighted equally within origins, rather than by the number of guts examined; the Tunapuna LU site had no available gut-content samples and was therefore not included in the calculation of origin-level mean occurrence frequencies.*

**Table S20. HU–LU differences in reconstructed dietary and somatic stoichiometry, diet-to-body indices, and elemental concentrations.**

| Response | Source | Unit | HU–LU [95% CrI] | Pr(direction) |
|---|---|---|---|---|
| C:N | Diet | Relative % | +42.7 [-9.0, 132.6] | 0.940 |
| C:N | Soma | Relative % | -4.2 [-12.4, 5.8] | 0.102 |
| C:N | Diet-to-body index | Relative % | +48.6 [-5.9, 145.1] | 0.957 |
| C:P | Diet | Relative % | -41.3 [-69.7, 10.5] | 0.046 |
| C:P | Soma | Relative % | -16.6 [-32.8, 3.8] | 0.043 |
| C:P | Diet-to-body index | Relative % | -29.6 [-64.9, 37.8] | 0.145 |
| N:P | Diet | Relative % | -59.2 [-81.8, -10.0] | 0.013 |
| N:P | Soma | Relative % | -12.8 [-29.8, 10.7] | 0.090 |
| N:P | Diet-to-body index | Relative % | -53.4 [-79.8, 6.5] | 0.035 |
| %C | Diet | Percentage points | -7.31 [-19.54, 5.40] | 0.125 |
| %C | Soma | Percentage points | -2.46 [-5.51, 0.56] | 0.038 |
| %N | Diet | Percentage points | -2.21 [-4.55, 0.33] | 0.044 |
| %N | Soma | Percentage points | -0.19 [-0.82, 0.44] | 0.171 |
| %P | Diet | Percentage points | +0.044 [-0.042, 0.161] | 0.857 |
| %P | Soma | Percentage points | +0.446 [-0.323, 1.205] | 0.911 |

*Values are posterior medians [95% credible intervals] for the high-minus-low urbanization (HU–LU) contrast. Relative % denotes 100 × (HU/LU − 1) for molar ratios and diet-to-body indices; percentage points denote differences in elemental concentrations (% dry mass). The diet-to-body index is the reconstructed dietary molar ratio divided by its mass-adjusted somatic counterpart. Pr(HU > LU) is the posterior probability that the contrast is positive (for negative contrasts, the probability of a decrease is 1 − this value).*
